# Synergistic targeting of cancer cells through simultaneous inhibition of key metabolic enzymes

**DOI:** 10.1101/2025.05.02.651820

**Authors:** Jan Dreute, Julia Stengel, Jonas Becher, David v. d. Borre, Maximilian Pfisterer, Marek Bartkuhn, Vanessa M. Beutgen, Benardina Ndreshkjana, Ulrich Gärtner, Johannes Graumann, Michael Huck, Stephan Klatt, Chloe Leff, Henner F. Farin, Andrea Nist, Roland Schmitz, Thorsten Stiewe, Julia Teply-Szymanski, Jochen Wilhelm, Alfredo Cabrera-Orefice, M. Lienhard Schmitz

## Abstract

As cancer cell specific rewiring of metabolic networks creates potential therapeutic opportunities, we conducted a synthetic lethal screen utilizing inhibitors of metabolic pathways. Simultaneous administration of (R)-GNE-140 and BMS-986205 (Linrodostat) preferentially halted proliferation of ovarian cancer cells, but not of their non-oncogenically transformed progenitor cells. While (R)-GNE-140 inhibits lactate dehydrogenase (LDH)A/B and thus effective glycolysis, BMS-986205, in addition to its known inhibitory activity on Indoleamine 2,3-dioxygenase (IDO1), also restricts oxidative phosphorylation (OXPHOS), as revealed here. BMS-986205, which is being tested in multiple Phase III clinical trials, inhibits the ubiquinone reduction site of respiratory complex I and thus compromises mitochondrial ATP production. The energetic catastrophe caused by simultaneous interference with glycolysis and OXPHOS resulted in either cell death or the induction of senescence in tumor cells, with the latter being eliminated by senolytics. The frequent synergy observed with combined inhibitor treatment was comprehensively confirmed through testing on tumor cell lines from the DepMap panel and on human colorectal cancer organoids. These experiments revealed highly synergistic activity of the compounds in a third of the tested tumor cell lines, correlating with alterations in genes with known roles in metabolic regulation and demonstrating the therapeutic potential of metabolic intervention.

## Introduction

The various hallmarks of cancer include significant alterations in metabolism. This allows tumors to meet their increased demand for building blocks to enable rapid proliferation ^1,2^. Tumor cell metabolites can also exhibit non-metabolic functions by regulating epigenetic changes upon supply of donor groups for acetylation (Acetyl-CoA) and methylation (S-adenosylmethionine). In addition, metabolites such as α-ketoglutarate serve as cofactors for dioxygenases ^3^, while fumarate bonds the antioxidant glutathione to amplify ROS-dependent signaling ^4^.

Metabolic reprogramming of cancer cells can be achieved in different ways. On the one hand, genetic alterations can be causative for metabolic shifts as described for isocitrate dehydrogenase 1 (IDH1), where the mutated enzyme converts α-ketoglutarate to 2-hydroxyglutarate ^5^. In addition, genes encoding metabolic enzymes such as phosphoglycerate dehydrogenase (*PHGDH*) can be amplified, leading to increased biosynthesis of serine, which is essential for proliferation of breast cancer and melanomas ^6,7^. On the other hand, metabolic reprogramming of cancer cells often occurs independently of genetic mutations in metabolic enzymes, arising instead as a consequence of oncogene expression. In this context, MYC confers glutamine dependence, which requires enhanced uptake of this amino acid and additionally promotes aerobic glycolysis ^8^. Oncogenic mutations of KRAS lead to increased uptake of glucose and channeling of its intermediates into the pentose phosphate pathway ^9^.

Changes in cancer metabolism were first described a century ago ^10^ and are frequently used for diagnosis of tumor cells using positron emission tomography (PET) imaging ^11^. Recent progress allows to determine the metabolic fingerprints of single circulating tumor cells to predict their distinct metastatic potential ^12^. The clinical relevance of therapeutic modulation of metabolic pathways is supported by meta-analyses of patients taking approved metabolic drugs such as statins or metformin for the treatment of hypercholesterolemia or type 2 diabetes. If these patients additionally developed tumors, the intake of statins or metformin resulted in improved survival rates and lower recurrence rates of various cancers such as ovarian cancer and breast cancer ^13,14^. Metabolic drugs have a range of benefits that justify their consideration in complex cancer therapies. Resistance to conventional tumor-targeting drugs such as cisplatin often creates metabolic vulnerabilities that are sensitive to treatment with metabolic inhibitors ^15,16^. Furthermore, their limited toxicity reduces the development of secondary malignancies that can occur later in life as a consequence of tumor therapy ^17^. Metabolic inhibitors are also used to reprogram the tumor microenvironment (TME), which can either restrain or support tumor proliferation ^18^. A range of therapies for the treatment of cancer are based on the therapeutic modulation of metabolic pathways. The administration of asparaginase leads to the cleavage of extracellular asparagine and is an FDA-approved approach to limit proliferation of acute lymphoblastic leukemia and lymphoblastic lymphoma, which both depend on the uptake of this amino acid ^19^. Interference with extracellular metabolites is also the rationale for phase III clinical trials of IDO1 (indoleamine 2,3-dioxygenase 1) inhibitors. This enzyme metabolizes tryptophan into the immunosuppressive metabolite kynurenine, which inhibits the therapeutically desired infiltration of tumors with cytotoxic CD8^+^ T cells ^20^.

The TME has been investigated in various suitable models including ovarian cancer ^21^. The vast majority of ovarian cancer patients are diagnosed with high-grade serous ovarian cancer (HGSOC), which typically originate from fallopian tube secretory epithelial cells. This disease is the deadliest gynecologic malignancy worldwide, with a 5-year survival rate of only about 35% ^22,23^. The frequent development of drug resistance in HGSOC prompts the need for innovative treatment options, such as the use of drug combinations targeting different pathways and reducing compensatory survival mechanisms ^24^. Furthermore, combining drugs may allow for lowering of individual dosages and reducing potential side effects. As the concept of synthetic lethality has also been applied to metabolic drugs ^25^, we set out to investigate the effects of combining metabolic inhibitors in ovarian cancer, a disease characterized by highly deregulated metabolism and supported by lipid transfer from neighboring adipocytes ^26,27^. For this, we compared immortalized fallopian tube secretory epithelial cells with their oncogenically transformed derivatives.

The synthetic lethal screen with metabolic inhibitors identified preferential suppression of tumor cell proliferation by a combination of (R)-GNE-140 (an inhibitor of the lactate dehydrogenases (LDH) A/B) and the IDO1 inhibitor BMS-986205, which is currently tested in clinical trials. The synthetic lethality of BMS-986205 was not attributable to IDO1 inhibition, but rather to a previously unknown off-target effect, identified here as the inhibition of complex I of the respiratory chain. The resulting energy shortage caused tumor cell senescence and allowed their elimination by senolytic drugs. The highly synergistic mechanism of action of both inhibitors was also observed in a large panel of tumor cells and patient-derived cancer organoids.

## Results

### Synthetic lethal targeting of metabolic pathways in ovarian cancer cells

To allow a direct side-by-side comparison between non-oncogenically transformed control cells and HGSOC tumor cells, a genetically defined model system was created. Since MYC and KRAS-mediated signaling is frequently deregulated in HGSOC ^28^, we generated a tumor model by oncogenic transformation of immortalized fallopian tube secretory epithelial cells (iFTSEC), which constitute the origin of HGSOC ^29^. These cells were transduced to stably express c-MYC together with hyperactive KRAS^G12V^, as schematically shown in Fig. 1A. The oncogene-expressing cells showed a significantly higher proliferation rate as compared to iFTSEC cells transduced with the empty expression vector (EV) (Fig. 1A). The KRAS^G12V^/MYC expressing tumor cells showed further characteristic features of cancer cells ^30,31^, such as a decreased cell size and cytoskeletal remodeling (Fig. 1B, Extended Data Fig. S1A), increased migration (Extended Data Fig. S1B, C) and trans-well invasion (Extended Data Fig. S1D, E). Consequently, only the KRAS^G12V^/MYC tumor cells had the ability to grow independently of anchorage, as revealed by soft agar colony formation assays (Fig. 1C). The KRAS^G12V^/MYC tumor cells also displayed the “Warburg effect”, as revealed by a higher rate of glycolysis quantified by Seahorse metabolic flux experiments (Fig. 1D, Extended Data Fig. S1F). In contrast, the generation of ATP within the mitochondria through OXPHOS remained unchanged (Extended Data Fig. S1G, H). Given the metabolic alterations exhibited by KRAS^G12V^/MYC expressing tumor cells, we aimed to exploit these metabolic differences as an entry point for selective targeting of cancer cells. Based on a number of criteria including target selectivity, potency, pharmacokinetics and dynamics, a literature search identified a range of suitable candidate inhibitors, as specified in suppl. Table 1.

**Fig. 1.**
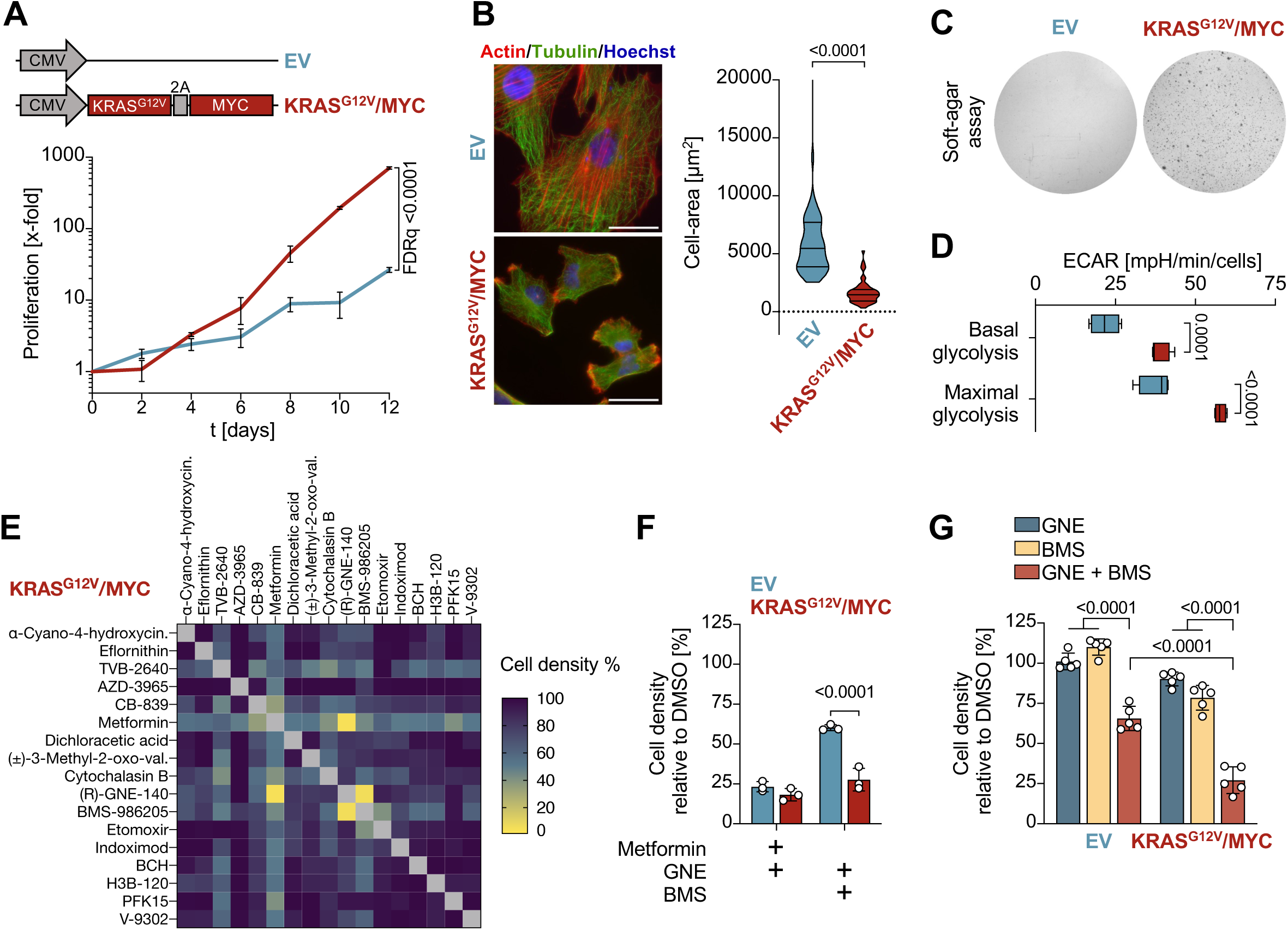
Synthetic lethality screen using engineered ovarian cancer cells. **(A)** Upper: immortalized fallopian tube secretory epithelial cells (iFTSEC) were transduced with a lentivirus encoding KRAS^G12V^ and MYC oncogenes separated by the porcine teschovirus-1 2A (P2A) sequence to allow for polycistronic gene expression or the empty vector as a control. Lower: three months after transduction and selection, proliferation was analyzed for 12 days. Shown are mean ± SD, normalized to t = 0 days, *n* = 3, multiple t test, FDR *q* values. **(B)** Left: KRAS^G12V^/MYC cells and EV control cells were stained for Actin and Tubulin and examined for cytoskeletal architecture and cell area by immunofluorescence microscopy. Scale bars = 50 µm. Right: Cell-area quantification of 51 cells, violin plots are shown and statistics are presented as two-tailed, unpaired t-test. **(C)** 2 × 10^4^ of the indicated cells were seeded in soft-agar and stained with crystal violet after 9 days of growth. **(D)** Extracellular acidification rate (ECAR) of the indicated cells was analyzed by Seahorse metabolic flux analysis, maximal ECAR was determined in the presence of oligomycin (2 µM). Shown are mean ± SD, *n* = 4, two-way ANOVA with Šídák’s multiple comparisons test. **(E)** Sublethal concentrations of the indicated metabolic inhibitors (suppl. Table 1) were added to KRAS^G12V^/MYC cells in 96-well plates. After 3 days, cell densities were determined by staining with crystal-violet and quantification in a plate reader. **(F)** The indicated combinations of Metformin, (R)-GNE-140 and BMS-986205 were added to the cells. After 3 days, cell densities were quantified by crystal-violet staining, data are shown as mean ± SD, *n* = 3, two-way ANOVA with Šídák’s multiple comparisons test. **(G)** KRAS^G12V^/MYC and EV control cells were treated with GNE (7.5 µM) and/or BMS (6 µM) for 3 days followed by determination of cell densities by crystal violet staining. Shown are mean ± SD, *n* = 5, two-way ANOVA with Dunnett’s and Šídák’s multiple comparisons test.

Sublethal concentrations of each inhibitor were determined, followed by a synthetic lethal screen in KRAS^G12V^/MYC tumor cells using non-toxic pairwise combinations of different inhibitors. Only two drug combinations suppressed proliferation of KRAS^G12V^/MYC tumor cells, namely (R)-GNE-140 in combination with either Metformin or BMS-986205 (Fig. 1E). To investigate the tumor cell specificity of these inhibitor combinations, their cytostatic effects were also tested on the immortalized EV cells. These experiments showed that only the combination of (R)-GNE-140 and BMS-986205 preferentially targeted the oncogene-transformed cancer cells (Fig. 1F). While (R)-GNE-140 (from hereon referred to as GNE) inhibits LDHA/B ^32^, BMS-986205 (from here on referred to as BMS, also known as Linrodostat) is known to interfere with tryptophan catabolism upon inhibition of indolamin 2,3-dioxygenase (IDO1) ^33^. Both drugs individually had only limited effects on KRAS^G12V/^MYC cancer cells, while a combination strongly impaired their proliferation (Fig. 1G).

### Drug synergy in human models

To test whether the effects of GNE and BMS are also seen in other cell systems, 19 genetically different cell lines were exposed to this drug combination. The analysis revealed a spectrum of responses, which were classified into three categories: high, medium and non-synergistically responding cells. (Fig. 2A). These data show that a combination of GNE and BMS can act in synergy to inhibit the proliferation of several cancer cell lines without displaying a general cytotoxicity. As cancer cell lines frequently exhibit genetic instability and accumulate a multitude of secondary mutations ^34^, we assessed the potentially synergistic effects of the combination therapy in primary non-cancerous cells and also in more complex human tumor models. As a model for immortalized, yet not entirely transformed human cells, we used *in vitro* cultivated germinal center B cells derived from cancer-free tonsil tissue that constitutively expressed MYC and BCL2 or BCL6 and BCL2, and subsequently cultured with feeder cells secreting growth-promoting cytokines (Extended Data Fig. S2A) ^35^. Treatment of these germinal center B cells with GNE/BMS did not show synergistic cell killing to the extent that was observed for tumor cells (Fig. 2B, Extended Data Fig. S2B). The drugs were then applied to KRAS^G12V^/MYC cancer cells that were grown as spheroids in soft agar, as such models replicate many aspects of the 3D structure and tumor environment in the body ^36^. The effectiveness and synergistic behavior of the combination therapy was also seen in KRAS^G12V^/MYC cancer cells grown as three-dimensional colonies (Fig. 2C). Subsequently we interrogated the effects of GNE/BMS treatment on patient-derived colorectal cancer organoids which retain important characteristics of primary tumors and adequately represent the inter-patient heterogeneity (Extended Data Fig. S2C) ^37^. Administration of GNE and BMS to tumor organoids revealed a broad spectrum of responses, ranging from non- to high-synergy (Fig. 2D, E, Extended Data Fig. S2D), reflecting the frequently observed heterogeneity in tumor cell sensitivity ^38^. GNE/BMS concentrations of all tested cell lines and organoids can be found in suppl. Table 2.

**Fig. 2.**
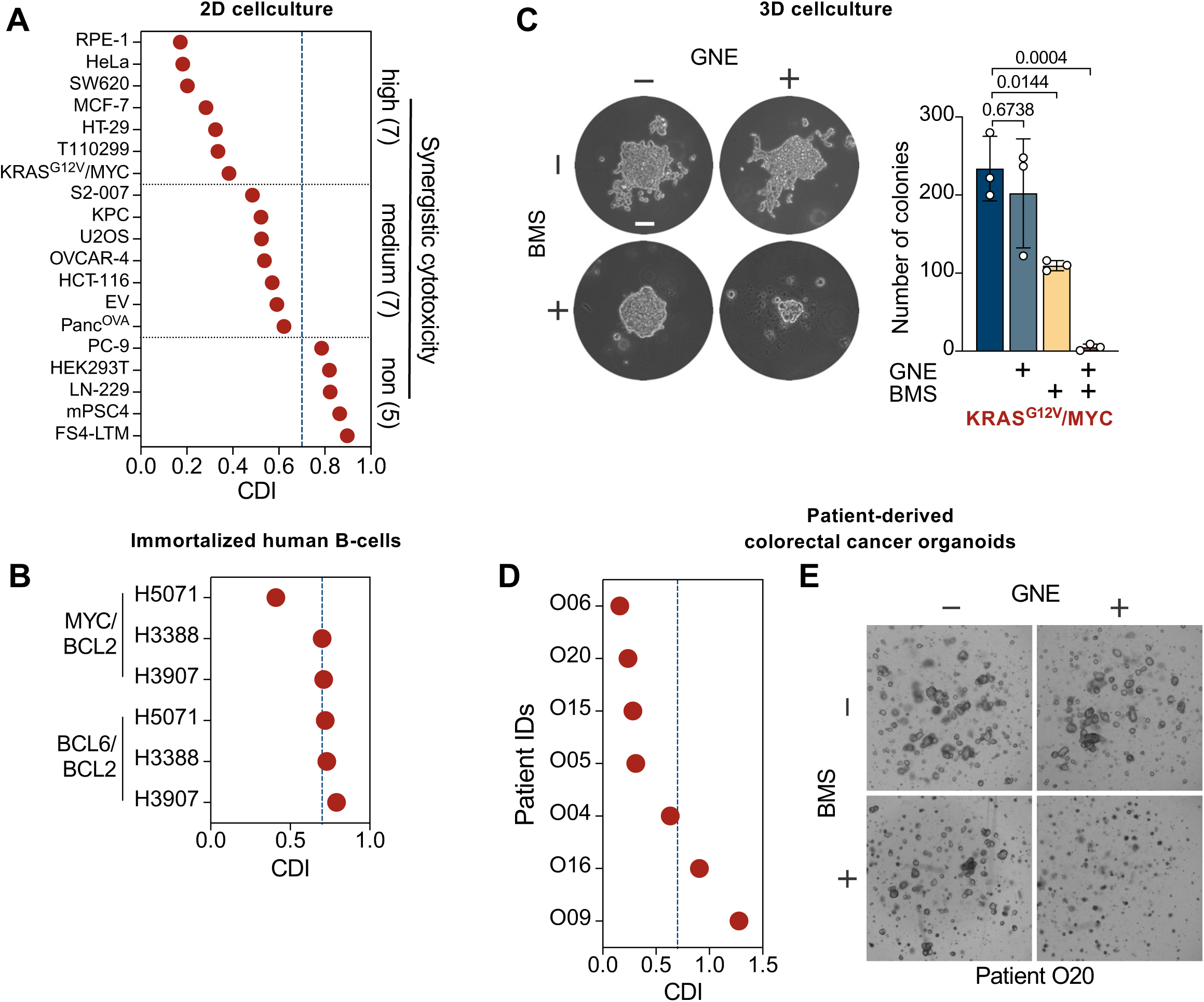
GNE/BMS synergy in different cell systems. **(A)** The indicated cell lines were treated with GNE and BMS (suppl. Table 2) either alone or in combination and cell densities were quantified by crystal violet staining. Synergistic inhibition of cell proliferation was calculated by determination of the Coefficient of Drug Interaction (CDI). CDI values ≥ 0.7 (indicated by a blue-dotted line) were considered to lack synergistic activity, while cells showing CDI values ≤ 0.4 show high synergy. **(B)** Primary germinal center B cells were transduced with MYC-BCL2 or BCL6-BCL2 and treated with sublethal concentrations of GNE and BMS (suppl. Table 2). After 7 days, cell number was determined and the average CDI score calculated, *n* = 4. **(C)** Left: KRAS^G12V^/MYC cells were grown in soft agar for 5 days to allow colony formation, followed by administration of (R)-GNE-140 and BMS-986205, which was repeated one week later. After 12 days, cells were stained with crystal violet. Scale bar = 30 µm. Right: Quantification of soft-agar colony formation experiments. Shown are mean ± SD, *n* = 3, two-way ANOVA with Dunnett’s multiple comparisons test. **(D)** Patient-derived colorectal cancer organoids were transduced with a lentivirus encoding Luciferase2-P2A-EGFP and selected with puromycin to enable stable expression. Organoids were harvested, singularized enzymatically and grown in 96-well round-bottom plates to organoids. Following treatment with GNE and/or BMS for 6 days (concentrations listed in suppl. Table 2), with a repeated treatment after 3 days, organoids were analyzed by microscopy and cell viability was assessed using the ONE-GloEX assay. The average CDI scores of different tumor organoids are shown, *n* = 2. **(E)** The morphology of organoids from patient O20 responding to GNE/BMS treatment is shown as an example.

### Combination treatment with GNE and BMS induces senescence in ovarian cancer cells

As KRAS^G12V^/MYC cancer cells respond to GNE/BMS combination treatment and allow comparison with their non-cancerous progenitor cells, follow-up experiments were conducted using this cell model for ovarian carcinoma and standard concentrations of GNE (7.5 μM) and BMS (6 μM). We performed RNA-seq experiments to study the effects of GNE/BMS on gene expression. Combination treatment of KRAS^G12V^/MYC cells for 4 hours resulted in expression changes of 1070 genes (log_2_FC ≥ 1, ≤ −1; FDR ≤ 0.05) (Fig. 3A, Extended Data Fig. S3A). To identify pathways associated with the early response of KRAS^G12V^/MYC cells to combination treatment, gene set enrichment analysis (GSEA) using the KEGG database was performed. GSEA revealed significant downregulation of genes annotated to cell cycle and DNA replication pathways, consistent with the observed anti-proliferative activity of GNE/BMS. Furthermore, GSEA revealed enrichment of genes annotated to pathways involved in post-translational modifications, signaling and extracellular matrix (Extended Data Fig. S3B). A longitudinal analysis of gene expression over 5 days post combination treatment revealed a large set of genes upregulated over time (cluster 1), while another set of genes showed time-dependent decrease of RNA abundance (cluster 2) (Fig. 3B, Extended Data Fig. S3C, D). GSEA using the KEGG (Fig. 3C) and Reactome databases (Extended Data Fig. S3E) assigned many of the downregulated differentially expressed genes (DEGs) from cluster 2 to regulators of cell survival and proliferation, while the upregulated transcripts contain many cytokines and chemokines. As the upregulated cluster 1 genes resemble a proinflammatory signature characteristic for a senescence-associated secretory phenotype (SASP) ^39^, we examined the GNE/BMS-regulated secretome in KRAS^G12V^/MYC cells by Olink^®^ Explore primer extension assays (PEA)^®^ (Extended Data Fig. S3F). These experiments revealed increased levels of 564 proteins (log_2_FC ≥ 1, ≤ −1; FDR ≤ 0.05) including the BMS target IDO1 after 4 days of treatment (Fig. 3D, Extended Data Fig. S3G). KEGG enrichment analysis of the secreted factors revealed that a considerable proportion of the identified proteins are inflammatory cytokines and matrix remodelers typical of SASP secretomes ^40^ (Fig. 3E).

**Fig. 3.**
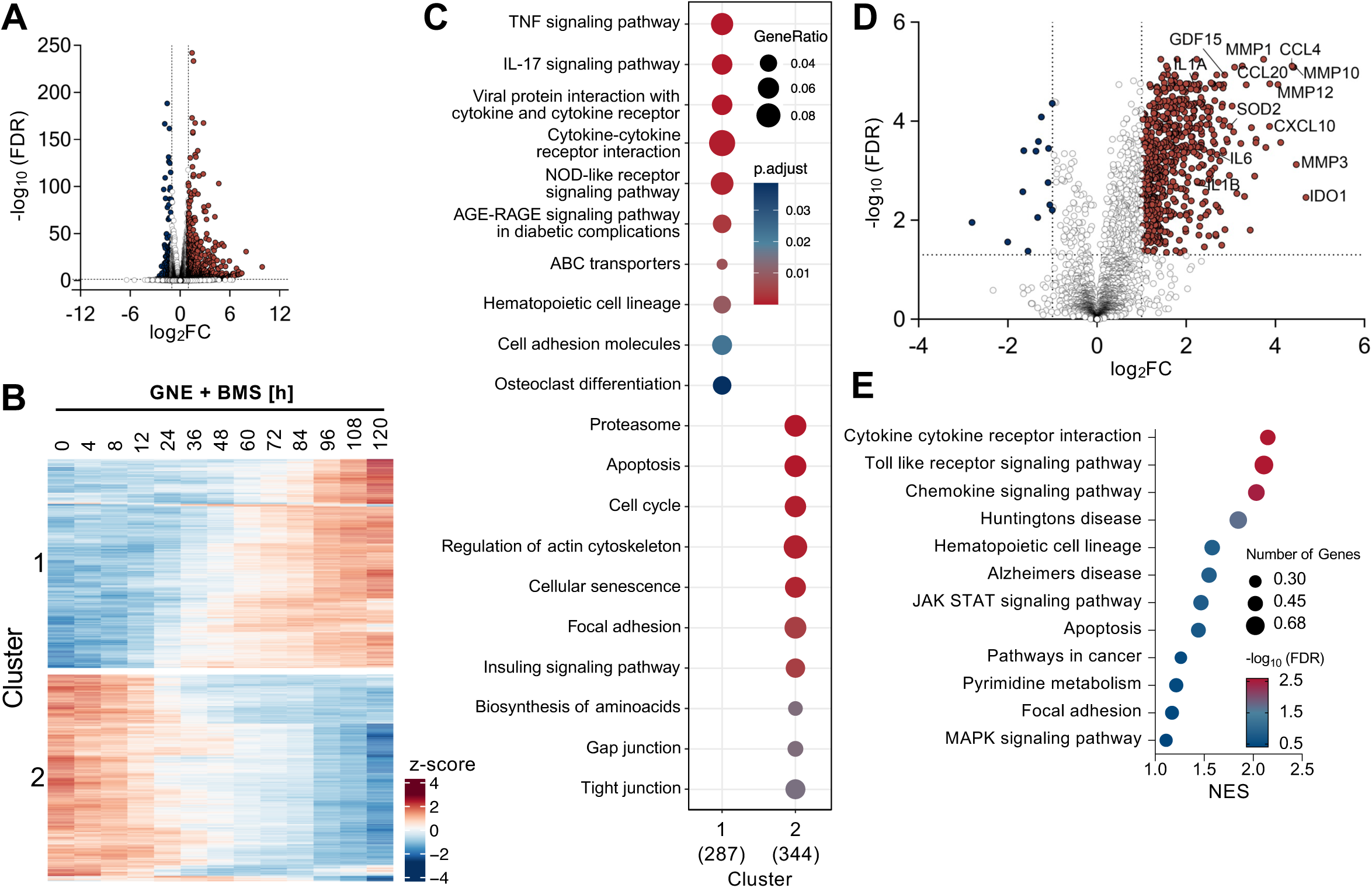
GNE and BMS combination treatment induces the senescence-associated secretory phenotype in KRAS^G12V^/MYC cancer cells. **(A)** KRAS^G12V^/MYC cells were treated for 4 hours with GNE and BMS or an adequate DMSO control, followed by analysis of gene expression by RNA-seq. A volcano plot is shown, *n* = 3. **(B)** KRAS^G12V^/MYC cells underwent GNE and BMS combination treatment for specified time periods, followed by RNA-seq. The top 1500 transcripts exhibiting the highest 3^rd^ order polynomial fits over time were k-means clustered (cluster 1 = upregulated over time, cluster 2 = downregulated over time). **(C)** The two different clusters were subjected to GSEA analysis using the KEGG database (KEGG disease database excluded). **(D)** KRAS^G12V^/MYC cells were treated for 4 days with GNE/BMS or DMSO vehicle control and secreted factors were quantified by Olink^®^. A volcano plot is shown, *n* = 4. **(E)** Differentially secreted proteins (log_2_FC ≥ 1, ≤ −1; FDR ≤ 0.05) were subjected to GSEA using the KEGG database.

To investigate whether GNE/BMS treatment induces senescence, the activity of senescence-associated ß-galactosidase (SA-ß-gal) was measured. The activity of this marker enzyme was significantly increased in KRAS^G12V^/MYC cells upon GNE/BMS combination treatment, as determined by brightfield microscopy (Fig. 4A, Extended Data Fig. S4A). Time-resolved analysis of SA-ß-gal activity showed significant induction already 2 days after treatment with maximal activity after 5 days, preferentially in KRAS^G12V^/MYC cells (Fig. 4B). As an increased cell size promotes senescence ^41^, we also determined the impact of both compounds on cell morphology. Microscopic analysis of GNE/BMS-treated cells showed that KRAS^G12V^/MYC cancer cells reacted with the formation of slender cell extensions and an increased size (Fig. 4C). Combination treatment triggered further hallmarks of senescence in KRAS^G12V^/MYC cells ^42^, such as increased concentrations of mitochondria-derived reactive oxygen species (mtROS), mainly superoxide anions (Fig. 4D). This elevation in ROS was accompanied by increased DNA damage, as detected by phosphorylation of the histone variant H2A.X (Fig. 4E, Extended Data Fig. S4B). Furthermore, GNE/BMS treatment led to an enrichment of KRAS^G12V^/MYC cells in the G2/M phase of the cell cycle and reduced incorporation of Bromodeoxyuridine (BrdU) (Fig. 4F, Extended Data Fig. S4C). Consistent with published data ^43,44^, the senescent cells showed reduced gene expression of the clinically relevant proliferation markers *MKI67*, *PCNA* and of the nuclear lamina protein lamin B1 (*LMNB1*) (Extended Data Fig. S4D). This reduction was accompanied by nuclear dysmorphology (Fig. 4G, H), which may contribute to the extensive number of dysregulated genes observed in the RNA-seq studies. We proceeded to explore whether the induction of senescence by GNE/BMS could open a new therapeutic opportunity to eradicate the senescent tumor cells using a senolytic drug ^45–47^. For this, cells were treated for 2 days with control conditions or the GNE/BMS combination to induce senescence, followed by application of Dasatinib for an additional 2 days (Fig. 4I). The senolytic compound Dasatinib, which suppresses anti-apoptotic signaling by interfering with the activity of several protein kinases ^48^, efficiently and selectively eliminated GNE/BMS-treated senescent KRAS^G12V^/MYC cancer cells, but not the control cells (Fig. 4J, K). Efficient elimination of senescent KRAS^G12V^/MYC cells by Dasatinib was also recapitulated in colony formation assays (Fig. 4L). This senolytic compound reprogrammed the cells from senescence to apoptosis, as shown by the cleavage and thus activation of caspase-3 (Fig. 4M).

**Fig. 4.**
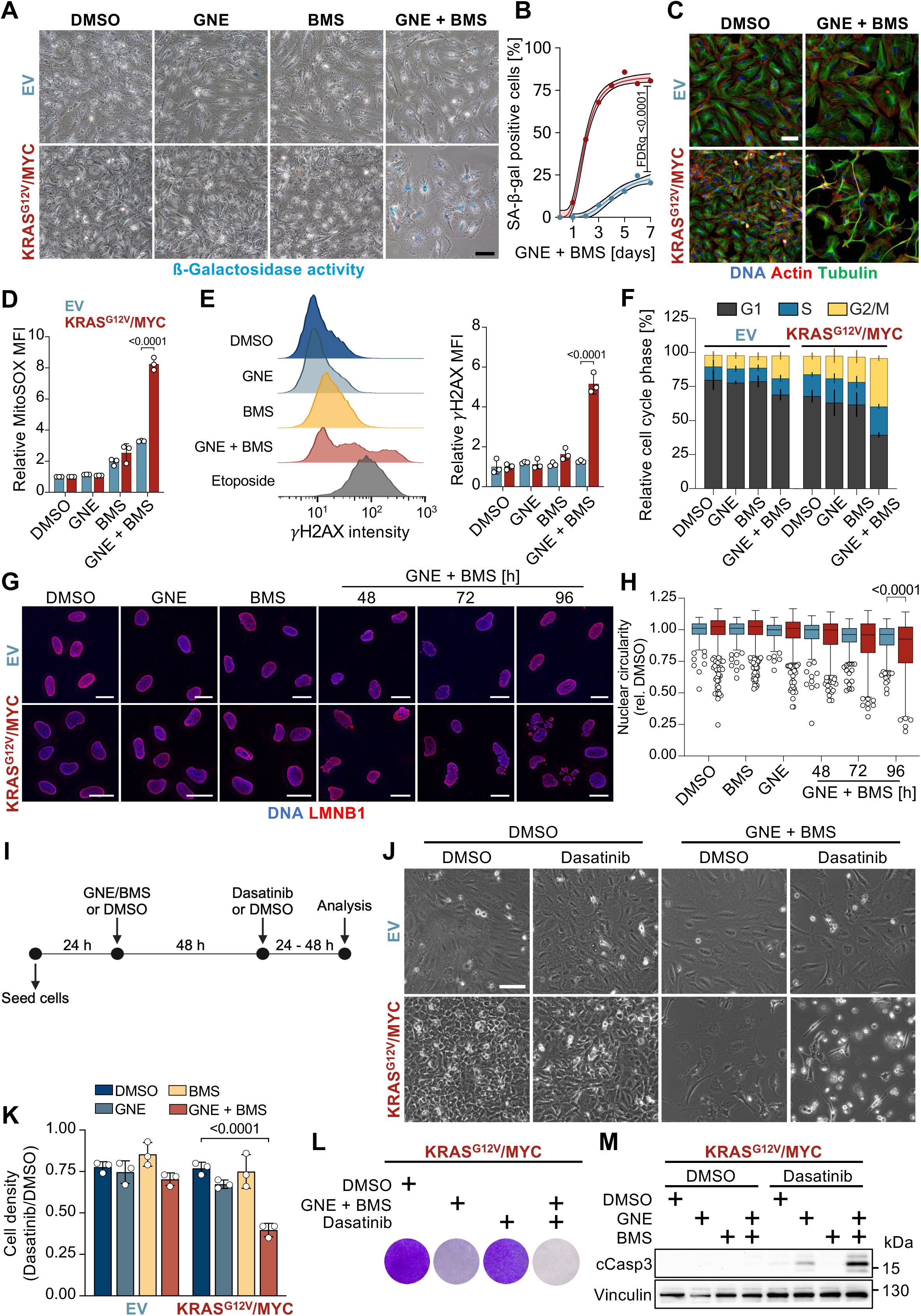
GNE/BMS-induced senescence enables therapeutic opportunities. **(A)** The indicated EV control and KRAS^G12V^/MYC cancer cells were treated for 3 days as shown, followed by determination of SA-ß-gal activity. A representative experiment is shown. Scale bar = 100 µm. **(B)** Quantification of SA-ß-gal positive cells over the course of 7 days after GNE and BMS combination treatment. Shown area mean and 95% confidence interval, *n* = 3, multiple t test, FDR *q* values. **(C)** The indicated cells were treated for 3 days with GNE and/or BMS, followed by analysis of the cytoskeletal components Actin and Tubulin via immunofluorescence. Scale bar = 100 µm. **(D)** Cells were treated for 2 days with GNE/BMS and loaded for 10 minutes with 5 µM MitoSOX^TM^ to detect mtROS by flow-cytometry. Shown are median fluorescence intensities ± SD normalized to DMSO, *n* = 3, two-way ANOVA with Šídák’s multiple comparisons test. **(E)** KRAS^G12V^/MYC cancer cells were investigated for the occurrence of DNA damage upon 2 days of treatment, as determined by flow-cytometric quantification of γH2AX. Etoposide treatment for 1 day (10 µM) was used as a positive control. Shown are histograms (left) of KRAS^G12V^/MYC cells and mean fluorescence intensities ± SD normalized to DMSO (right), *n* = 3, two-way ANOVA with Šídák’s multiple comparisons test. **(F)** Cells were treated as shown and the distribution of cell cycle phases was determined by flow-cytometric quantification of propidium iodide-stained cells, *n* = 3. **(G)** KRAS^G12V^/MYC cancer cells and EV controls were treated as shown, followed by Lamin B1 immuno-staining and confocal-fluorescence imaging. Scale bar = 25 µm. **(H)** The Lamin B1 signal was used for segmentation and calculation of nuclear circularity. Shown are boxplots with Tukey whiskers, *n* = 3, one-way ANOVA with Šídák’s multiple comparisons test. Outlier values are presented as circles. **(I)** Schematic workflow representation of senolytic compound addition. EV control and KRAS^G12V^/MYC cell lines were exposed to GNE/BMS for 2 days, followed by addition of Dasatinib (50 nM) or Mock (DMSO) for another 2 days. **(J)** Cells were microscopically inspected and representative brightfield images of indicated conditions are shown. Scale bar = 100 µm. **(K)** Cell density quantification using crystal violet staining. Dasatinib treated conditions were normalized to Mock treated conditions to show the Dasatinib-specific effect. Shown are mean ± SD, *n* = 3, two-way ANOVA with Dunnet’s multiple comparisons test. **(L)** KRAS^G12V^/MYC cells were treated as in (A), followed by washing away of non-adherent cells and staining of attached cells using crystal violet, a representative result is shown. **(M)** The indicated cell lines were treated for 2 days with GNE/BMS and then for another day with Dasatinib (50 nM). Cell extracts were analyzed by Western blotting for cleavage of caspase 3 as a marker for apoptosis, *n* = 3.

### The synthetic lethality caused by BMS-986205/Linrodostat relies on its effect on mitochondrial structure and function

Are the cytostatic effects of GNE and BMS attributable to inhibition of their known targets? To address this question, LDHA, LDHB and IDO1 were either downregulated with siRNAs or treated with other inhibitors targeting LDHA/B (GSK-2837808A) or IDO1 (Epacadostat). These experiments failed to recapitulate the strong synergistic effects seen with GNE and BMS on cell survival (Extended Data Fig. S5A-C), suggesting the occurrence of an off-target effect. As elevated ROS levels often derive from mitochondria ^49^, we investigated these organelles in further detail. To visualize mitochondria in intact cells, KRAS^G12V^/MYC and EV control cells were transduced to allow stable expression of EGFP in fusion with COX8, a component of respiratory complex IV. Cells with mitochondrial COX8-EGFP were treated with the drugs alone or in combination. Only the combination of both compounds led to a significant increase in the number of mitochondria per cell, as seen by confocal fluorescence microscopy and mitochondrial 3D-rendering, selectively in the KRAS^G12V^/MYC cells (Fig. 5A, B, Extended Data Fig. S5D). The increased number of mitochondria upon GNE/BMS treatment was also observed by flow-cytometric quantification of MitoTracker labeled mitochondria (Extended Data Fig. S5E) and was accompanied by a decrease of mitochondrial branch length (Fig. 5C). Transmission electron microscopy (TEM) revealed that treatment with BMS was sufficient to reduce folding of the inner mitochondrial membrane (Fig. 5D). We evaluated the impact of both compounds on mitochondrial respiration and glycolysis with seahorse metabolic flux experiments. In line with the BMS-induced changes in mitochondrial structure, this compound led to an almost complete shut-down of OXPHOS (Fig. 5E, Extended Data Fig. S5F-H). Quantification of extracellular acidification revealed the expected downregulation upon LDHA/B inhibition by GNE (Fig. 5F, Extended Data Fig. S5I, J). Interestingly, the respective inhibitory effect on OXPHOS or glycolysis was accompanied by a slight compensatory induction of the other, unhindered ATP production pathway. The energy map shown in Fig. 5G visualizes that GNE causes a compensatory upregulation of aerobic ATP production by the respiratory chain, while BMS causes a compensatory shift to glycolytic energy production. Combination treatment blocks ATP production by glycolysis and OXPHOS and thus shifts the cells into an energy quiescent state. A treatment-specific difference was uncovered upon determination of the extracellular pH, which is lowered by lactate excretion. As BMS leads to a shut-down of OXPHOS, the compensatory upregulation of glycolysis resulted in lactate excretion and acidification of the cell-culture medium, while administration of GNE resulted in the basification of the medium in a plethora of cell lines (Fig. 5H), which is in line with its known function as an LDH inhibitor ^50^. These dynamic pH changes were significantly more pronounced in the tumor cells compared to the EV cells, consistent with their higher metabolic activity.

**Fig. 5.**
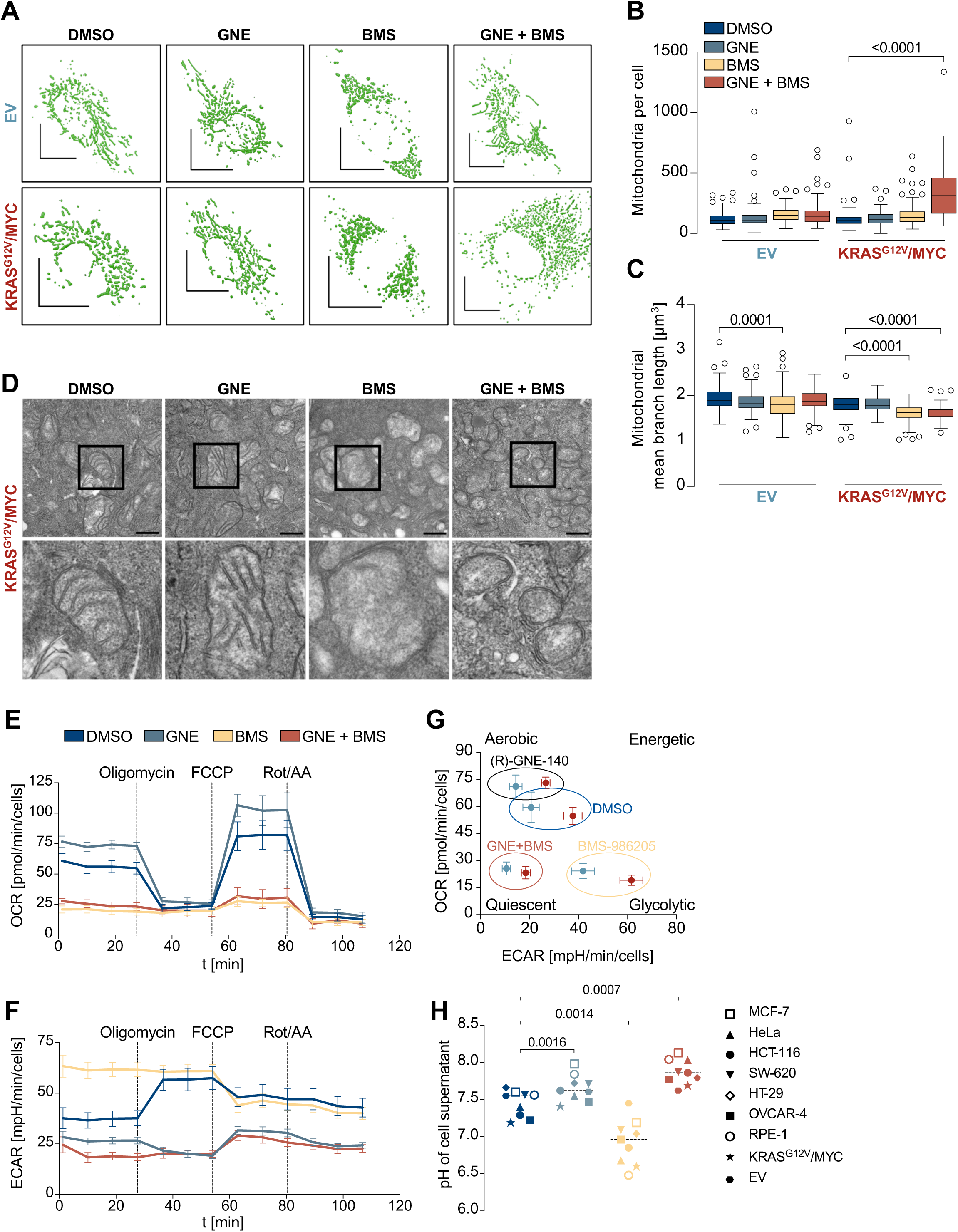
Mitochondrial effects of BMS. **(A)** The indicated EV control and KRAS^G12V^/MYC cancer cells were transduced to stably express COX8-EGFP, enabling visualization of mitochondria. Cells were treated for 2 days with indicated conditions, followed by fixation and confocal imaging. High-resolution 3D rendered mitochondria of representative cells are displayed. Scale bar = 10 µm. **(B)** The number of mitochondria per cell and **(C)** mitochondrial mean branch length was quantified, *n* = 3, one-way ANOVA with Šídák’s multiple comparisons test. **(D)** KRAS^G12V^/MYC cells were treated as indicated, followed by analysis of mitochondrial structure via transmission electron microscopy. Scale bar = 500 nm. The rectangular selections are shown in the lower row at a higher magnification. **(E)** KRAS^G12V^/MYC cancer cells were treated with GNE and/or BMS for 4 hours, followed by Seahorse metabolic flux analysis. Oligomycin, FCCP and Rotenone/Antimycin A were injected into the wells at the indicated timepoints, followed by quantification of the oxygen consumption rate (OCR). **(F)** The experiment was performed as in (E) for quantification of the extracellular acidification rate (ECAR). Seahorse experiments show mean ± SD, *n* = 4. **(G)** Energy map summarizing the Seahorse experiments and annotating the different treatments to metabolic states (Aerobic, Glycolytic, Energetic or Quiescent). **(H)** The indicated cell lines were treated for 4 days with GNE and/or BMS and the extracellular pH was determined. Statistical analysis was done using two-way ANOVA with Dunnett’s multiple comparisons test.

### BMS-986205/Linrodostat: a novel inhibitor of complex I

To directly test a possible effect of BMS on the activity of the respiratory chain complexes, KRAS^G12V^/MYC and HEK293T cells were fractionated to enrich mitochondria. Bovine heart mitochondria, the standard model system for the studies of mitochondrial function ^51^, were isolated and purified using differential centrifugation and a sucrose gradient. Isolated mitochondria were then subjected to a standardized protocol allowing assessment of enzymatic activities of respiratory chain complexes I-IV ^52^. The addition of BMS caused a selective and dose-dependent decrease of complex I activity in mitochondria from KRAS^G12V^/MYC, HEK293T cells and bovine heart (Fig. 6A, B, C).

**Fig. 6.**
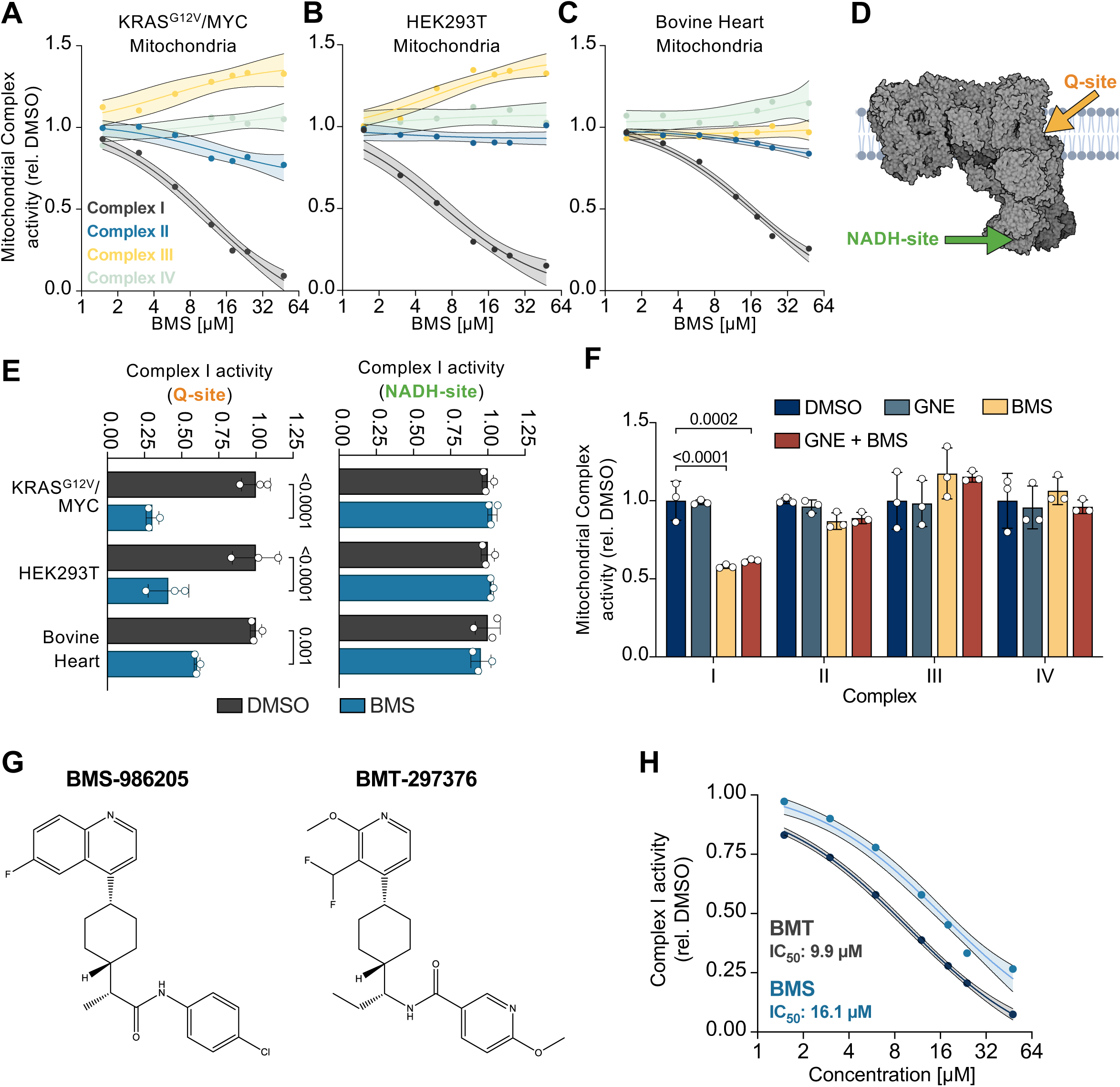
BMS inhibits the Q site of complex I. **(A, B, C)** Mitochondria were isolated from KRAS^G12V^/MYC, HEK293T cells and bovine heart tissue. Respiratory chain complex-specific activity assays were performed in the presence of the indicated BMS concentrations or with the highest percentage of DMSO (0.48 % (v/v)) as a control, and analysed spectrophotometrically. *n* = 3 for complex I, *n* = 2 for complexes II-IV. **(D)** Schematic representation of the ubiquinone (Q) and NADH binding sites within complex I. **(E)** Isolated mitochondria were tested for the activities of the Q and NADH sites of complex I in the presence of BMS (24 µM) or DMSO (0.36 % (v/v)). Mean ± SD are shown, *n* = 3, two-way ANOVA with Šídák’s multiple comparisons test. **(F)** The experiment was performed as in (A-C) in the presence of GNE (7.5 µM) and/or BMS (6 µM). Mean ± SD are shown, *n* = 3, two-way ANOVA with Dunnett’s multiple comparisons test. **(G)** Chemical structures of BMS-986205 and BMT-297376. **(H)** The indicated concentrations of the two compounds were added to mitochondria and complex I activity was determined. Mean and 95% confidence interval are shown, *n* = 2.

Most known inhibitors of complex I target the ubiquinone binding site (Q-site) or the NADH site, which are located at different positions within complex I (Fig. 6D). To distinguish between these two sites, spectrophotometric assays for NADH oxidation and ubiquinone reduction were performed. BMS specifically interfered with the ubiquinone reduction activity in mitochondrial fractions of different origins, without displaying any impact on the electron transfer at the NADH site (Fig. 6E). Further experiments demonstrated that BMS exhibited pronounced selective activity against complex I, whereas GNE did not contribute to the inhibitory effects (Fig. 6F). Together, these data show that BMS, which has already been used in multiple phase III clinical trials (NCT03661320, NCT03329846), is not only a potent IDO1 inhibitor, but also displays a significant off-target effect by inhibiting the Q-site of complex I and thereby the main entry of electrons into the respiratory chain. This finding has broad implications for the further use and safety profile of BMS, making it relevant to investigate whether structurally related IDO1 inhibitors also share this off-target effect. Experiments addressing this question showed that also the IDO1 inhibitor BMT-297376 (a next-generation BMS-986205, see Fig. 6G) displayed selective inhibitory activity for complex I (Fig. 6H), urging to reconsider the specificity of this class of BMS-related inhibitory molecules.

### GNE/BMS leads to exhaustion of energy carriers and building blocks for proliferation

To investigate the metabolic consequences of GNE/BMS combination treatment, a targeted metabolomic analysis was performed for amines, nucleotides and building blocks of the central carbon metabolism. In KRAS^G12V^/MYC cells exposed to GNE and/or BMS for 4 hours, single treatment induced changes in only a few metabolites, while the combination treatment led to stronger deregulation, primarily resulting in downregulation of metabolic compounds (Fig. 7A, B). Combination treatment resulted in the decrease of the energy carriers ATP and phosphocreatine (Fig. 7B, C) and consequently activated the energy-sensing AMPK pathway, as observed by phosphorylation of the acetyl-CoA carboxylase (ACC) at serine 79 (Fig. 7D). Activation of AMPK activity was paralleled by inhibition of mTOR activity, as reflected by decreased phosphorylation of its direct target eukaryotic translation initiation factor 4E-binding protein 1 (4E-BP1) at threonine 37 and 46 and serine 65 ^53^. Furthermore, the combination treatment induced significant deregulation of 35 metabolites, including the exhaustion of nucleotides and acetyl-CoA, one of the central hubs in carbon metabolism (Fig. 7E) ^54^. Enrichment analysis of deregulated metabolic pathways revealed the overrepresentation of the Warburg effect (glycolysis), but also of pathways contributing to redox homeostasis (pentose phosphate pathway, glutathione metabolism) and proliferation (purine and pyrimidine metabolism) (Fig. 7F).

**Fig. 7.**
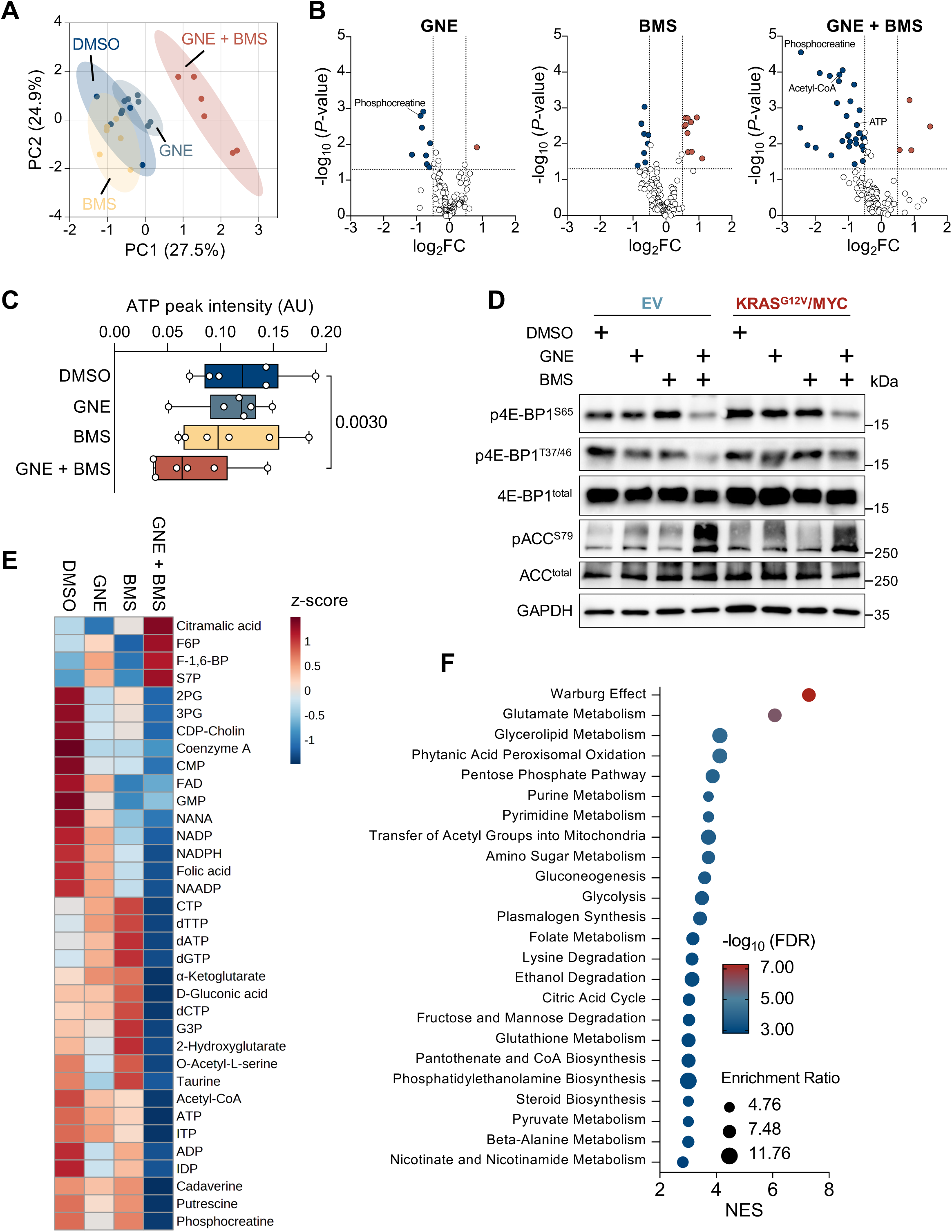
GNE/BMS treatment leads to metabolic deregulation. **(A)** KRAS^G12V^/MYC cells received GNE and/or BMS for 4 hours, followed by targeted metabolomic analysis using LC-MS. Shown is a partial least squares discriminant analysis of the specified samples using MetaboAnalyst 5.0. Shaded areas display the 95% confidence interval. **(B)** Volcano plots illustrate the metabolic alterations induced by the indicated inhibitors in comparison to DMSO treatment, *n* = 6. **(C)** Boxplot showing ATP peak intensity, *n* = 6, two-tailed paired t test. **(D)** The indicated cells were treated with BMS and/or GNE for 4 hours and equal amounts of proteins were analyzed by Western blotting. Specific antibodies detect the expression and phosphorylation of proteins reflecting the activity of mTOR kinase (4E-BP1 phosphorylation) and AMPK (ACC phosphorylation). The positions of molecular weight markers are shown. **(E)** Significantly deregulated metabolites upon GNE/BMS combination treatment (log_2_FC ≥ 0.5, ≤ −0.5; *P* ≤ 0.05) are shown in a heatmap depicting z-scores per metabolite. **(F)** The significantly deregulated metabolites were subjected to enrichment analysis and mapped against the SMPDB database using MetaboAnalyst 5.0. Only those pathways which yielded more than four metabolites were considered.

### Molecular pathways underlying the GNE/BMS synergy

To evaluate the combination therapy in different tumor entities we subsequently screened a large panel of tumor cell lines of different origins and with known molecular signatures. A total of 102 human cancer cell lines were examined for their synergistic response to GNE/BMS combination treatment, including 86 deriving from solid tumors and 16 hematologic cancer cell lines. A strongly synergistic effect (Bliss score > 20, dotted line) was observed for 70 cell lines (Fig. 8A). The cell lines with the most synergistic behavior originated from the pancreas and ovary (Fig. 8B), supporting the suitability of the ovarian cancer model mainly used in this study. Since the organ-type alone cannot explain the sensitivity towards the GNE/BMS combination, we correlated the responsiveness to genomic alterations using a subset of the most commonly occurring and intensively studied cancer genes (suppl. Table 3). This analysis revealed a significant link between synergistic behavior and mutations of genes encoding the low-density lipoprotein receptor-related protein 1B (*LRP1B*), the acetyl transferase p300 (*EP300*) and the transcriptional corepressor 1 (*RB1*) (Fig. 8C, suppl. Table 4). Interestingly, the products of these mutated genes have known roles in regulating metabolic cues. ^55–57^. Further statistical analyses were performed using basal gene expression levels of 95 Oncolines^®^ cell lines retrieved from the DepMap database (version 23Q4). Correlation analysis between cell line sensitivity to GNE/BMS combination and basal gene expression levels of 19,080 genes revealed that high expression of mRNAs encoding *TP53* (wildtype and mutant) may serve as a marker of cell line sensitivity (Pearson’s r = 0.33, *P* value = 0.00096) (Fig. 8D). A potential marker of cell line resistance to GNE/BMS is high expression of the PI3 kinase catalytic subunit (*PIK3CA*) (Pearson’s r = −0.24, *P* value = 0.017). In this context it is interesting to note that the senolytic drug Dasatinib used in this study for elimination of GNE/BMS treated cells (see Fig. 4K), also interferes with PI3K signaling ^58^. Since it is unlikely that the expression level of a single gene alone determines responsiveness to GNE/BMS treatment, we tried to identify gene sets that correlate with sensitivity or resistance. For this, a pre-ranked gene list, according to the full list of Pearson correlations, was subjected to GSEA using the KEGG database. The analysis revealed an enrichment of gene sets associated with GNE/BMS treatment sensitivity, encompassing the spliceosome, cell cycle, and related processes, including DNA replication and mismatch repair pathways (Fig. 8E). It is noteworthy, that these synergy-associated gene sets overlap strongly with the gene sets found to be downregulated upon 4 hours of combination treatment identified in RNA-seq experiments (Extended Data Fig. 3B). Interestingly, the seven most negatively enriched gene sets, thus indicating treatment resistance, are comprised of metabolic pathways with implications in energy homeostasis (linoleic-, retinoic acid metabolism) or drug-detoxification (CYP450-family).

**Fig. 8.**
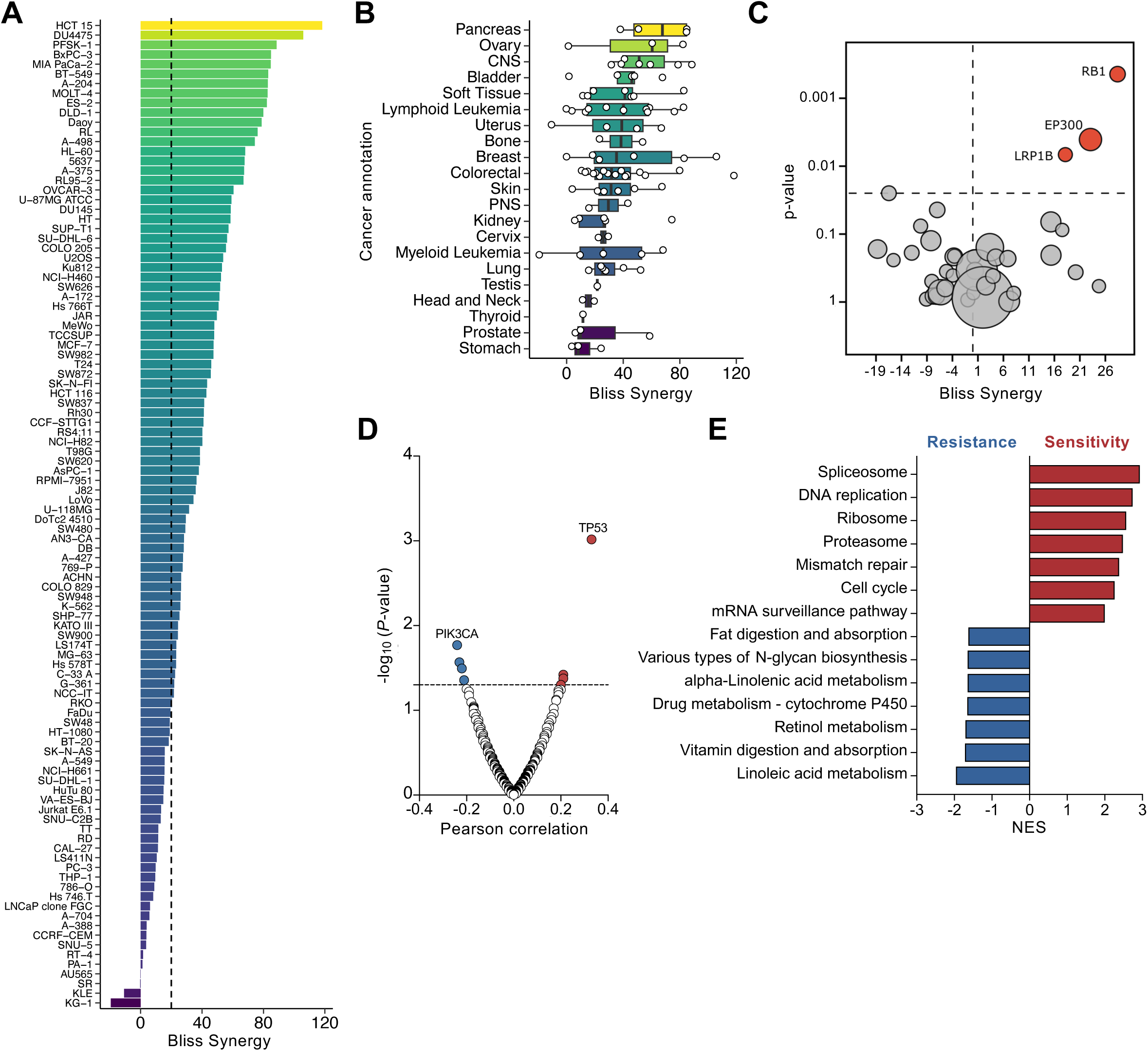
Determinants of GNE/BMS synergy. After determination of sublethal drug concentrations of GNE and BMS, the indicated cell lines were treated for 3 days using the previously determined, cell line-specific, fixed sublethal concentration of BMS in combination with the full dose-titration of GNE. The effect of GNE/BMS combination on cell viability was determined by measuring the intracellular ATP content by ATP Lite^TM^. **(A)** GNE/BMS synergy was calculated using the Bliss independence model. The dotted line indicates the threshold of high Bliss synergy (> 20). **(B)** Cell lines were grouped according to their tissue of origin and presented as boxplots. **(C)** Volcano plot showing the synergy behavior of combination treatment in relation to mutations in 38 known tumor-relevant genes (suppl. Table 3 and 4). **(D)** Volcano plot of Pearson correlations between GNE/BMS Bliss synergy scores and basal expression levels of 19,080 genes of 95 cell lines. Shown is a subset of cancer relevant genes. **(E)** The pre-ranked Pearson correlation list of 19,080 genes was subjected to GSEA using the KEGG database as the reference set. Depicted are the top 7 enriched pathways associated with either GNE/BMS resistance or sensitivity.

## Discussion

This study demonstrates that joint targeting of two critical metabolic pathways can result in pronounced synergistic effects on tumor cell viability. Such an approach has several advantages: It allows interference with metabolic routes despite the inherent plasticity of tumor cell metabolism. Furthermore, combination therapy can limit the mechanisms by which a tumor can escape therapeutic intervention. For example, OXPHOS inhibition often leads to upregulation of glycolysis, thus the addition of LDH inhibition can counteract this compensatory metabolic adaptation. In addition, combination therapy prevents the development of resistance frequently seen with the use of LDH inhibitors ^32^. This combination therapy may also reshape the TME by reducing the secretion of immunosuppressive and tumor-promoting metabolites such as lactate and kynurenine, as a consequence of LDH and IDO1 inhibition, respectively ^59,60^.

BMS-986205 and its derivative BMT-297376 are reported to inhibit indoleamine 2,3-dioxygenase (IDO1) ^33,61^ and, as revealed in this study, also the Q-site in complex I. It is quite possible that this activity may indirectly cause the observed architectural changes in mitochondria, as comparable alterations have been observed in the presence of other Q-site inhibitors, such as rotenone ^62^. Inhibition of complex I frequently results in reduced ATP production and increased ROS levels ^63^. These processes, along with other metabolic alterations, caused by inhibition of IDO1 and LDH, along with changes of mitochondria and their biochemical pathways, can then trigger response programs such as apoptosis or senescence. Currently, the mechanisms distinguishing between these different cell fates remains unclear. However, numerous tumors become senescent as (i) they show metabolic adaptations to cope with cell increased proliferation, and (ii) the mere expression of oncogenes can be sufficient to prime cells for senescence ^64^.

Mitochondrial dysfunction is both a cause and consequence of cellular senescence, playing a key role in various feedback loops that initiate and maintain the senescence phenotype. ^65^. The GNE/BMS treatment results in reduced ATP levels, which increase AMPK activity. This, in turn, is known to contribute to p53-independent cell cycle arrest via FOXO3/Gadd45a ^66,67^. Off-target effects of clinically used tumor drugs on mitochondria are not without precedent, as exemplified by doxorubicin, which in addition to inhibiting its target topoisomerase II, also interferes with OXPHOS through various mechanisms ^68,69^. Similarly, cisplatin preferentially binds mitochondrial DNA and kills tumor cells dependent on the production of mitochondrial ROS ^70,71^. In addition, all-trans-retinoic acid, which is used for differentiation treatment of patients with acute promyelocytic leukemia, inhibits the mitochondrial adenine nucleotide translocase and triggers the intrinsic apoptosis pathway ^72^.

Since many anti-tumor drugs have side effects that directly or indirectly affect mitochondrial function ^73^, these unintended effects may even be relevant for the effectiveness of the therapy. Tumor-promoting or antagonizing functions of mitochondria do not only rely on their contribution to a number of metabolic pathways, but also on their relevance for various cell death pathways including apoptosis and ferroptosis ^74,75^. In addition, mitochondria serve as sensors and integrators for distinct stress-induced signaling pathways ^76^. Mitochondria can even foster tumor cell metastasis, as revealed by replacing mitochondrial DNA of a poorly metastatic mouse tumor cell line with mutated mitochondrial DNA from a metastatic cell line ^77^. The importance of mitochondria for the cytotoxic effects of BMS is underscored by the observation that metformin, the other substance that showed synergistic tumor toxicity in combination with GNE, also interferes with respiratory complex I activity ^78–80^.

The synergism between GNE and BMS was not observable in all cell systems tested, and several tumor cell lines did not respond to combination treatment. We assume that these cell lines satisfy their ATP demands by alternative pathways such as basal glycolysis, which can proceed in the absence of LDHA/B activity along the Embden-Meyerhof-Parnas pathway ^81^. In addition, cells may bypass complex I by utilizing other electron donors (e.g., succinate, fatty acids, ketone bodies, etc.) and thus allowing continued ATP production via OXPHOS, albeit with reduced efficiency ^82^.

However, further studies are needed to elucidate the underlying mechanisms defining the responsiveness to GNE/BMS treatment. Among the frequently mutated candidates in synergistically reacting cells is the p300 protein, which utilizes acetyl-CoA as a co-substrate, a compound among the key proxies reflecting the overall metabolic state of the cell ^83^. Impaired function of p300 and further candidate proteins including LRP1B and RB1 could possibly skew metabolic pathways to become vulnerable towards GNE/BMS treatment. Interestingly, we found that cells showing low response or resistance to GNE/BMS display a high basal mRNA expression of the catalytic subunit of PI3K. This protein is known to play a central role in promoting pro-survival and therapy resistance, raising the possibility of pharmacologically manipulating cell responsiveness through PI3K inhibition ^84,85^. The identification of cell properties that define a synergistic response to GNE/BMS combination treatment is of paramount importance and could potentially enable precision targeting of responsive tumor cells through the use of prodrugs or tailored drug delivery strategies ^86,87^. In such a scenario, targeting of tumor cells by specific delivery of GNE/BMS together with senolytic drugs could enable selective elimination of tumor cells.

## Methods

### Cell culture

Immortalized fallopian tube secretory epithelial cells (iFTSEC) ^88^ were purchased from abm^®^ (FT240, cat. #T0762-GVO-ABM). Cells were grown in DMEM/F-12 GlutaMAX supplemented with 10% fetal bovine serum, 100 U/ml penicillin and 100 µg/ml streptomycin at 37°C with 5% CO_2_. Empty vector and KRAS^G12V^/MYC cell lines were generated using the 3^rd^ generation lentiviral packaging system (pMDLg/pRRE, pRSV-rev, pHCMV-VSV-G). pLenti CMV Blast empty (w263-1) was a gift from Eric Campeau & Paul Kaufman (Addgene plasmid #17486; RRID:Addgene_17486). KRAS^G12V^ and c-MYC sequences were codon-optimized and synthesized by BioCat and cloned into the donor-vector. Lentiviral particles were generated in HEK293T cells, according to manufacturer’s instructions. Supernatants were collected and centrifuged followed by filtration. Filtrated supernatant was mixed with 6 μg/ml polybrene (Sigma, #H9268) and used 1:5 with complete medium for transduction of iFTSECs. After 16 hours of transduction, medium was replaced with fresh complete medium. For the next 3 months, cells were selected using 7.5 µg/ml Blasticidin, phenotypically monitored and subsequently tested for cancerous transformation. For GFP-labeling of mitochondria, empty vector and KRAS^G12V^/MYC cell lines were transduced to express COX8-GFP (Addgene plasmid #44385; RRID:Addgene_44385, from Pantelis Tsoulfas). GFP positive cells were enriched by FACS sorting. Cell lines allowing for short tandem repeat (STR) profiling (RPE-1, HeLa, SW620, MCF-7, HT-29, S2-007, U2OS, OVCAR4, HCT-116, PC9, HEK293T, LN229) were STR authenticated. STR-authentication for FT240 was conducted by the supplier abm^®^. Cell lines were regularly tested for Mycoplasma contamination (Minerva biolabs, Venor^®^GeM Classic).

### Analysis of germinal center B cells

Tonsil tissue was provided by the department of Otolaryngology, Head and Neck Surgergy of the University Hospital of Giessen and Marburg. Primary human germinal center B cells were isolated from tonsillar tissue with prior written informed consent from patients/parents/guardians as previously described ^89^. The study received approval from the institutional ethics committee of the Justus Liebig University, Giessen (approval number: AZ37/20). GC B cells were co-cultured on irradiated YK6-FDC (follicular dendritic cells)-like feeder cells expressing CD40L and IL21 in advanced RPMI 1640 medium supplemented with 10% fetal bovine serum, 100 U/ml penicillin and 100 µg/ml streptomycin at 37°C with 5% CO2. Prolonged GC B cell culture was enabled by retroviral transduction with either MSCV-BCL6-t2a-BCL2 or MSCV-MYC-t2A-BCL2 oncogenic backbone, a gift from Daniel Hodson (Addgene plasmid #135305 and #135306, respectively; RRID:Addgene 135306). Cells were subcultured for a total of 10 days. The day before treatment, cells were seeded at 2 × 10^4^ cells per well in a 48 well plate followed by GNE/BMS mono and combination treatment. After 7 days of treatment, the cell number was determined using a Beckman-Coulter CytoFLEX S flow cytometer, gating for the lymphocyte population.

### Colorectal cancer organoid culture

Resection samples from patients with colorectal cancer were provided by the University Cancer Center Frankfurt (UCT). All materials were collected as part of the interdisciplinary Biobank and Database Frankfurt (iBDF), with prior written informed consent from patients. The study received approval from the institutional review board of the UCT and the Ethical Committee at the University Hospital Frankfurt (ethics vote: 4/09; project number: SGI-10-2022). Patient-derived organoids were cultured as described ^37^. Organoids were individually transduced with a Luciferase2-P2A-EGFP lentivirus as described ^90^ and stable expression was selected by expansion in the presence of 0.5 to 1 μg/mL puromycin. For drug testing, tumor organoid lines were seeded after enzymatic digest using Accutase (Thermo Fisher) and filtration (40 μm, Greiner) in 15 µL of 70% Cultrex UltiMatrix Reduced Growth Factor Basement Membrane Extract (Bio-Techne) in 96-well round-bottom plates (Sarstedt) in duplicates. Each well received 100 µL of medium containing advanced DMEM/F12 supplemented with 10 mM HEPES, 1× Glutamax, 1× penicillin/streptomycin, 2% B27, 12.5 mM N-acetylcysteine, 500 nM A83-01 (R&D Systems), 10 μM SB202190 (Sigma-Aldrich), 20% R-spondin 1 conditioned medium, 10% Noggin conditioned medium, 50 ng/mL human EGF (Peprotech), and Wnt surrogate (35 ng/mL, # N001-0.5 mg, ImmunoPrecise) for three days and then upon the drug administration reduced growth factor medium was used (advanced DMEM/F12 supplemented with 10 mM HEPES, 1× Glutamax and 1× penicillin/streptomycin). The plate was sealed with Breathe-easy membranes (Sigma-Aldrich, #Z380059) to prevent evaporation. On day 3, the medium was changed using an automatic multichannel pipette (Integra Mini-96), and the drug combination treatments were dispensed using a D300e digital dispenser (Tecan) in 7-point dilutions ranging from 0.5 to 50 µM. The DMSO content was normalized to the highest volume in all wells, ensuring it did not exceed 1% of the final volume. The cells were treated for 6 days in total, with a repeated treatment after 3 days. Morphological images (2.5X) of the entire 96-well plate were captured with a Cytation C10 confocal imaging reader (Agilent), and cell viability was assessed using the ONE-GloEX assay (Promega) according to the manufacturer’s instructions. Luminescence was measured with a SpectraMax iD3 Microplate Reader (Molecular Devices) and the raw data was normalized to the DMSO control.

### Analysis of cell migration and invasion

Scratch assay: 16 hours before initiating the scratch, complete cell medium was replaced with 1% FCS-containing medium to diminish cell proliferation. A vertical scratch of the cell monolayer was performed with a 200 µl tip. Subsequently, the medium was replaced to remove floating cells, and images of the same spot within each well were captured over the next 24 hours. Quantification of the scratch area was performed by the “Wound_healing_size_tool-3” plugin in FIJI ^91^. Transwell invasion: Cells were seeded into 6-well plates and cultured overnight. Following overnight culture, the medium was replaced with serum-free medium, and cells were starved for 24 hours. After trypsinization, cells were pelleted, resuspended in 1 ml serum-free medium containing CellTrace^TM^ Oregon Green^TM^ 488 (5 μM), and incubated for 30 minutes at 37°C. Simultaneously, 15 μl of type I Collagen Solution (bovine, 6 mg/ml) was added onto the transwell membrane, followed by a 45-minute incubation at 37°C. Coated surfaces were subsequently rinsed with PBS. After a 30-minute staining period, cells were washed twice with 1x PBS. 3000 cells were added to the upper compartment of the transwell chamber. The lower compartment was filled with 250 μl/well of serum-free media. Cell migration (in the absence of collagen) and invasion (in the presence of collagen) were allowed for 24 hours at 37°C. After 24 hours, the medium from both the upper and lower compartments was aspirated, and the upper compartment was cleaned with a cotton swab to remove non-migrated or non-invaded cells. Both chambers were rinsed once with 100 μl 1x PBS. The upper and lower compartments were then filled with 50 μl and 250 μl PBS, respectively. Images of the GFP^+^ cells were acquired using a Leica LAS X THUNDER microscope.

### Soft-agar colony formation assay

Soft-agar colony formation assay was performed in 6-well plates using noble-agar (Sigma, #A5431) prepared with 2x complete medium (2xFCS; 2xL-Glutamine). A 2 ml layer of 0.6% bottom-layer was poured into each well and solidified for 15 minutes at 4°C. Subsequently, 1 ml of 0.3% cell-layer containing 2 × 10^4^ cells was pipetted onto the bottom-layer. 500 μl growth medium was replenished biweekly using 2x complete medium. Cells were allowed to form visible colonies for 5 days before treatment. GNE and BMS mono- or combination treatments were applied for a total of 12 days, with drug replenishment once after 7 days.

### Determination of cell density by crystal violet

Cell density in response to treatment was measured using crystal violet assay ^92^. In brief, cells were seeded in 96-well plates (or other well-formats) overnight in duplicates. Cells were treated with inhibitors the next day and analyzed after 24 −72 hours of treatment as follows: Cells were washed twice with ice-cold PBS, placed on ice and fixed with ice-cold 100% Methanol for 10 minutes. After fixation, methanol was aspirated and cells were air-dried for 10 minutes at RT. Cells were incubated with staining solution (25% Methanol, 0.5% crystal violet in ddH_2_O) for 10 minutes. Excess staining solution was poured off and cells were washed three times with ddH_2_O. Cells were then air-dried at 37°C and subsequently de-stained for 30-60 minutes using 2% SDS in ddH_2_O. Absorbance was measured at 600 nm using a GloMax Discover^®^ plate reader (Promega, cat. no. GM3000). Background absorbance was measured from empty wells and subtracted from all values. Absorbance values were normalized to control cells which received adequate DMSO concentrations.

### ß-Galactosidase activity assay

Cells were washed two times with PBS and fixed in 2% formaldehyde, 0.2% glutaraldehyde in PBS for 5 minutes. X-gal staining solution (20 mM citric acid; 40 mM sodium hydrogen sulfate; 150 mM NaCl; 5 mM potassium ferrocyanide trihydrate; 5 mM Potassium ferricyanide; 2 mM MgCl2; 1 mg/ml X-gal in ddH_2_O) was added, 6-well plates were sealed with parafilm and incubated light-protected in an incubator overnight. Cells were washed with 100% methanol at room temperature. Images were acquired after several hours of light-protected drying.

### Immunofluorescence imaging

Experiments were performed using 18 mm glass-cover slips in 6-well plates. Cells were seeded at 50-70% confluency. Fixation was performed using 4% formaldehyde followed by blocking with 3% BSA-0.3% Triton-X-100 in PBS for 30 minutes. Between every step, cells were washed twice with 0.3% Triton-X-100 in PBS. Antibodies were diluted in PBS containing 0.3% BSA and 0.3% Triton-X-100 and incubation steps were carried out in a humidified chamber. Primary antibody incubation was conducted overnight at 4°C, while secondary antibody incubation occurred for 2 hours at room temperature. For BrdU staining, DNA strands were denatured for 1 hour at room temperature using 2 M HCL, prior to primary antibody incubation. The following primary antibodies were used (dilution, supplier, clone, cat#, RRID): BrdU (1:1000, Abcam, BU1/75 (ICR1), #ab74546, AB_1523225), LaminB1 (1:1000, Abcam, #ab16048, AB_443298), H2A.X Ser139 (1:1000, Millipore, JBW301, #05-636, AB_309864). Secondary antibodies (dilution, supplier, cat#): goat anti-rat IgG-Alexa Fluor 488 (1:1000, Dianova, #112-545-167), goat anti-rat IgG-Alexa Fluor 594 (1:1000, Dianova, #112-585-167), goat anti-rabbit IgG-Alexa Fluor 488 (1:1000, Dianova, #111-545-003), goat anti-rabbit IgG-Alexa Fluor 594 (1:1000, Dianova, #111-585-144), goat anti-mouse IgG-Alexa Fluor 488 (1:1000, Dianova, #115-545-003), goat anti-mouse IgG-Alexa Fluor 594 (1:1000, Dianova, #115-585-062). Cells were analyzed on an Eclipse TE2000-E inverted fluorescence microscope 350 (Nikon) equipped with a cooled pE-300 light source, an ORCA Spark CMOS camera (Hamamatsu), a Canon EOS 650D color camera and a T-RCP Controller (Nikon) using Nikon lenses (10X/0.3, 20X/0.4, 60X/1.4, 100X/1.4). Images were recorded with NIS Elements 3.10. Confocal images were acquired with an Aurox Unity spinning disk confocal microscope using the Aurox app.

### Western blotting

Cells were lysed in NP40 buffer (20 mM Tris/HCl, pH 7.5; 150 mM NaCl; 1 % (v/v) IGEPAL CA-630) with freshly added inhibitors (10 mM NaF; 0.5 mM Na_3_VO_4_; 1 mM PMSF; 10 μg/ml Aprotinin; 10 μg/ml Leupeptin). Protein yield was quantified using Pierce^TM^ BCA assay (ThermoFisher, #23227). For SDS-PAGE, 5-20 µg of total protein was used and semi-dry blotted on PVDF membranes. Blocking was performed in 5% nonfat dry milk or 5% bovine serum album (BSA) in in TBST (5 mM Tris, 15 mM NaCL, 0.1% Tween 20, pH 7.5). Primary antibody incubation was performed in 5% nonfat dry milk or BSA in TBST overnight at 4°C on a shaker. The following primary antibodies were used (Dilution, Supplier, Clone, cat#, RRID): ACC (1:1000, Cell Signaling, C83B10, #3676, AB_2219397), pACC Ser79 (1:1000, Cell Signaling, #3661, AB_330337), cleaved Caspase-3 Asp175 (1:1000, Cell Signaling, #9664, AB_2070042), GAPDH (1:200, DSHB, 2G7, #DSHB-hGAPDH-2G7), IDO1 (1:1000, Cell Signaling, D5J4E, #86630, AB_2636818), LDHA (1:1000, Cell Signaling, C4B5, #3582, AB_2066887), LDHB (1:5000, Proteintech, #14824-1-AP, AB_2134953), p4E-BP1 Ser65 (1:500, Santa Cruz, 62.Ser 65, #sc-293124, AB_2943677), p4E-BP1 Thr37/46 (1:1000, Cell signaling, 236B4, #2855, AB_560835), Vinculin (1:1000, Sigma, hVIN-1, # V9131, AB_477629). Secondary antibodies (Dilution, Supplier, cat#): goat anti-rat IgG-HRP (1:5000, Dianova, #112-035-143), goat anti-rabbit IgG-HRP (1:5000, Dianova, #111-035-144), goat anti-mouse IgG-HRP (1:5000, Dianova, #115-035-146). After washing and incubation with the appropriate horseradish peroxidase-coupled secondary antibodies, the antibody-bound proteins were detected using a Bio-Rad ChemiDoc imaging system.

### Analysis of mitochondrial morphology

COX8-GFP expressing cells were imaged on an Aurox Unity spinning-disk confocal microscope using a Nikon 60×/1.4 NA lens with a spatial resolution of 0.15 × 0.15 × 0.3 µm/px (XYZ). Images were processed using FIJI/ImageJ ^93^. Crosstalk between channels was corrected by linear unmixing. Background was removed by subtracting a minimum intensity projection from the Z-stack for each channel. Images were subsequently deconvolved using Microvolution with a theoretical PSF from PSFGenerator ^94^ using a Richards & Wolf model. Mitochondrial morphology was measured using the ImageJ plugin Mitochondria Analyzer ^95^ after optimization of thresholding parameters. Thresholded images of mitochondria were used to generate surfaces using ImageJ 3D Viewer and then rendered for visualization in Blender (https://www.blender.org). Analysis code is available under 10.5281/zenodo.14591823.

### Nuclear circularity analysis

Cells were stained for DNA (Hoechst33342) and Lamin B1 (1:1000, Abcam, #ab16048, AB_443298) and imaged on an Aurox Unity confocal microscope using a Nikon 60X/1.4NA lens at 0.15 x 0.15 x 0.3 µm/px resolution (XYZ). Briefly, nuclei were segmented from the Lamin B1 channel after background subtraction and BC adjustment by intensity-based thresholding. Cell doublets were manually removed and circularity was measured. Analysis code is available under 10.5281/zenodo.14591792.

### Transmission electron microscopy

For TEM analysis, cell pellets were fixed in 1.5 % glutaraldehyde and 2 % formaldehyde in 0.15 M HEPES/KOH (pH 7.4) and after washing post-fixed in a buffer containing 1% osmium tetroxide (Merck). After washing in distilled water, the samples were incubated overnight in 2 % aqueous uranyl acetate (Merck) at 4 °C, dehydrated in ethanol, and embedded in AGAR 100 (Agar Scientific Ltd, Essex, UK). Ultrathin sections were mounted on grids and analyzed using a transmission electron microscope (Zeiss EM 900EL, Carl Zeiss GmbH, Oberkochen, Germany) equipped with a slow-scan 2 K CCD camera (TRS, Tröndle, Moorenweis, Germany).

### Cell cycle analysis

Cells were fixed in on 70% EtOH and DNA was stained using propidium iodide in the presence of 200 µg/ml RNase A for 15 minutes at 37°C in the dark. DNA content was determined by flow cytometry.

### Determination of mtROS and mitochondrial abundance

Cells were trypsinized, transferred to Eppendorf tubes, and incubated with MitoSox Red (ThermoFisher, #M36008, 5 µM) for 10 minutes, followed by MitoTracker Deep Red FM (ThermoFisher, #M22426, 200 nM) for 15 minutes at 37°C in the dark. After incubation, cells were washed once with PBS and analyzed by flow cytometry.

### Seahorse metabolic flux analysis

For metabolic flux analysis, the Seahorse XF Cell MitoStress Test Kit (Agilent, cat. no. 103015-100) was used according to the manufacturer’s instructions. Cells were seeded in DMEM/F-12 complete medium in 96XF cell culture plates overnight. The next day, cells were treated with (R)-GNE-140 and/or BMS-986205 for 4 hours, followed by the MitoStress Test assay using Oligomycin (2 µM), FCCP (2 µM), and Antimycin A/Rotenone (0.5 µM). Seahorse measurements were obtained using the recommended Seahorse XF Cell MitoStress Test analysis, run on an XF Pro Analyzer. For in-well normalization of cell numbers, a second plate of cells was seeded and treated exactly the same way as the cells subjected to Seahorse analysis, with the exception that this normalization plate did not receive the MitoStress Test drugs (Oligomycin, FCCP, AntA/Rot). During the Seahorse analysis run, the normalization plate was subjected to crystal violet cell density quantification to control for GNE/BMS treatment and cell line-specific cell number variation.

### siRNA-mediated gene silencing

Cells were seeded at 40-60% confluence 1 day prior transfection. Target gene expression was silenced using small interfering RNAs (siRNAs) obtained from OriGene. Eukaryotic cells were transfected with Lipofectamine (ThermoFisher^®^) following the manufacturer’s instructions. In all study experiments, a concentration of 10-20 nM for a single siRNA was used. To increase the efficiency of gene silencing, two rounds of siRNA transfection were conducted. The following siRNAs were used (5’-3’): siIDO1-CGUAAGGUCUUGCCAAGAAAUAUTG, siLDHA-CUCCUGAAGUUAGAAAUAAGAAUGG, siLDHB-UUAUGAUGCAAUCAGGACUGUACUUGA.

### Library preparation and RNA sequencing (RNA-Seq)

For genome-wide analysis of gene expression, RNA sequencing libraries from isolated mRNA were generated and sequenced by the Institute for Lung Health (ILH) – Genomics and Bioinformatics – at the Justus-Liebig-University (JLU) Giessen (Germany). A total amount of 1000 ng of RNA per sample was used to enrich for polyadenylated mRNA using the NEBNext® Poly(A) mRNA Magnetic Isolation Module (New England BioLabs) followed by cDNA sequencing library preparation utilizing the NEBNext® Ultra™ II Directional RNA Library Prep Kit for Illumina® (New England BioLabs) according to the manufacturer’s instructions. After library quality control by capillary electrophoresis (4200 TapeStation, Agilent), cDNA libraries were sequenced on the Illumina NovaSeq 6000 platform generating 50 bp paired-end reads.

### RNA-seq data analysis

The Illumina software bcl2fastq (v2.19.0.316) was utilized for demultiplexing and generating FASTQ files. Initial processing of the sequencing read comprising quality control, filtering, trimming, alignment, and the creation of gene-specific count table was conducted using the nf-core RNA-seq v3.7 bioinformatics pipeline (^96^; NEXTFLOW version 23.04.03), with the homo sapiens hg38 genome and gene annotations sourced from Illumina’s iGenome repository (https://support.illumina.com/sequencing/sequencing_software/igenome.html). The pipeline ran with default parameter settings in docker mode. Subsequent analysis of the resulting count matrix, including raw read count normalization and differential gene expression detection, was performed in R using DESeq2 ^97^ with default settings. In case of the time series analysis, the limma package ^98^ was used to fit the data to a 3rd order polynomial after vst-transformation and scaling of the count matrix. PCAs were calculated based on the top 500 most highly variable genes. Pathway analysis was performed using the clusterProfiler package ^99^ against KEGG and Reactome databases. For the time series analysis, gene set enrichment analysis was performed for the contrasts of all individual time points versus the control sample. Heatmaps were created using the ComplexHeatmap package ^100^. In case of the time series analysis, the top 1500 genes fitted to the 3rd order polynomial where selected. The tree resulting from hierarchical clustering was cut into two groups (characterized by genes predominantly up-regulated and down-regulated over time). The KEGGREST (https://bioconductor.org/packages/KEGGREST; R package version 1.46.0) package was used for removal of disease terms from KEGG pathways. Raw sequencing data and derived gene expression matrix have been uploaded to NCBI’s Gene Expression Omnibus under accession number GSE285726. Analysis code is available under 10.5281/zenodo.14575774.

### Secretome analysis using Olink Explore 3072

KRAS^G12V^/MYC cells were treated with DMSO (0.135%) or (R)-GNE-140 and BMS-986205 combination for 4 days. The medium was harvested and used for further analysis. Affinity-based proteomic analysis was performed at the Core Facility Translational Proteomics (Philipps-University Marburg) using the Olink Explore 3072 platform, following the standard protocol (v1.5, 2022-12.21) and as described ^101^. Randomized samples were plated on a 96-well plate and processed in one batch. The generated libraries were sequenced by the Genomics Core Facility (Philipps-University Marburg) using Next-Generation Sequencing (NGS) on an Illumina NovaSeq6000 sequencer. The Olink Explore platform is based on proximity extension assay (PEA) technology ^102^ combined with a high-throughput NGS readout ^103^. This semi-quantitative platform reports protein levels as “Normalized Protein eXpression” (NPX), log_2_ scaled arbitrary units. Statistical analyses were performed with the *OlinkAnalyze* R package version 4.0.2. Analysis code and raw input files are available under 10.5281/zenodo.14591830.

### Targeted metabolome analysis

Amines were extracted by adding methanol extraction buffer (1 mM TCEP, 1mM ascorbic acid, 0.1% formic acid in 85:15 Methanol:H_2_O containing the internal standards of Homotaurine and Serotonin-d4, both at a final concentration of 0.001 mM) at a volume to cell pellet weight ratio of 10:1. The mixture was shortly vortexed, followed by 4 rounds of sonication. Protein fraction was removed via centrifugation. Supernatants were diluted with 2x volume of MilliQ water and frozen at −80°C following freeze-drying overnight. Dried samples were reconstituted in 80 μl boric acid buffer (200 mM boric acid in MilliQ water) followed by addition of 4 μl of 1N NaOH and 20 μl AQC reagent (6-aminoquinolyl-N-hydroxysuccinimidyl carbamate in 100% acetonitrile). Central carbon metabolites, nucleotides and nucleosides were extracted by addition of Trifluorethanol:H_2_O (1:1) and 10 minute incubation, followed by addition of MeOH:EtOH (1:1) for 10 minutes. After addition of ddH_2_O and incubation for 10 minutes, the samples were loaded onto Captiva EMR-Lipid Plates and pushed through. Captiva plates were washed with ddH_2_O:MeOH:EtOH (2:1:1) containing the internal standards citrate-1,5,13C2 and succinate-1,4-13C2. The combined filtrate was dried under nitrogen flow until solvents evaporated and subsequently shock frozen with liquid nitrogen and freeze-dried overnight. Dried samples were reconstituted with AcN and sonicated on ice for 45 seconds, followed by addition of ddH_2_O and sonication for 45 seconds on ice. Samples were centrifuged at 14000 rpm for 10 minutes at 4°C and the supernatants were transferred to MS glass vials and injected into the mass spectrometer (Agilent Technologies 6495C QQQ). Blank samples, quality controls and standard curve samples were injected. HPLC: 1290 Infinity II BIO (For Amines: Zorbax Extend C18 RRHD (2.1 x 150 mm, 1.8µ), Agilent; For Central Carbon, Nucleosides and Nucleotides: InfinityLab Poroshell 120 HLIC-Z (RRHD 2.1 x 150 mm, 2.7-Micron) PEEK Lined, Agilent). Peak-intensities (AU) were normalized to internal standards and the sample specific protein amount, as determined by Pierce^TM^ BCA assay (ThermoFisher, #23227). Normalized data were log_2_ transformed and subjected to downstream analyses as indicated, using MetaboAnalyst 5.0.

### Mitochondrial respiratory chain enzyme activity assays

#### Isolation of mitochondria from cancer cells

HEK293T or KRAS^G12V^/MYC cancer cells were seeded in T175 cell culture flasks and harvested at 95% confluency. Cells were trypsinised, washed and pelleted at 1000 x g for 5 min at 4°C and subsequently resuspended in SET buffer (0.25 M sucrose, 1 mM EDTA, 20 mM Tris-HCL, pH 7.5). Cells were disrupted by homogenization (Potter-Elvehjem tube) with 10 gentle strokes followed by pressing the suspension 5 times through a 23G syringe. All steps were performed on ice. The crude mitochondrial fraction was isolated using two centrifugation steps (1^st^: 600 x g, 10 min; 2^nd^: 12.000 x g, 10 min). Mitochondria were diluted to 1 mg*ml^-1^ protein in hypotonic 25 mM Tris-HCL, pH 7.5 followed by 2 freeze-thaw cycles to disrupt mitochondrial membranes. All enzymatic assays were performed at 37°C using a Tecan M200 Infinite spectrophotometer. All reactions were measured in 96-well plates at a final volume of 200 µl (path length ∼0.48 cm). Respiratory chain activity assays were performed as previously described ^52^. Sucrose gradient purified bovine heart mitochondria were isolated via differential centrifugation and purified via sucrose gradient. Enzyme activity was determined spectrophotometrically in all cases.

#### Complex I activity assay

Activity was determined by measuring NADH oxidation (ε_340nm_ = 6.2 mM^-1^*cm^-1^). Mitochondria from bovine heart (10 µg), HEK293T and KRAS^G12V^/MYC (50 µg) were resuspended in 25 mM potassium-phosphate buffer pH 7.5 and supplemented with 2 mg/ml fatty acid-free BSA and 20 µM oxidized cytochrome c. For the NADH-site assay (NADH:ferricyanide oxidoreductase activity), mitochondria were resuspended in 25 mM Tris-HCl, pH 8.5 and supplemented with 10 µM Rotenone, 2 µM Antimycin A and 1.2 mM Ferricyanide. For the Q-site assay (NADH:decylubiquinone oxidoreductase activity), mitochondria were resuspended in 25 mM potassium-phosphate buffer pH 7.5 and supplemented with 3 mg/ml fatty acid-free BSA and 70 µM decylubiquinone. The reaction was started upon the addition of 120 µM NADH. Complex I specific activity was determined by subtraction of the Rotenone (10 µM)-sensitive activity.

#### Complex II activity assay

Activity was determined by measuring 2,6-dichlorophenolindophenol (DCPIP) reduction (ε_600nm_ = 19.1 mM^-1*^cm^-1^). Mitochondria from bovine heart (5 µg), HEK293T and KRAS^G12V^/MYC (25 µg) were resuspended in 25 mM potassium-phosphate buffer pH 7.5 and supplemented with 1 mg/ml fatty acid-free BSA, 10 µM Rotenone, 2 µM Antimycin A, 20 mM succinic acid and 80 µM DCPIP. Succinic acid was added to the samples around 10 min before starting the assay to avoid competitive inhibition by endogenous oxaloacetate. The reaction was started upon addition of 70 µM decylubiquinone. Complex II specific activity was determined by subtraction of the malonate (10 mM)-sensitive activity.

#### Complex III activity assay

Activity was determined by measuring cytochrome *c* reduction (ε_550nm_ = 18.5 mM^-1*^cm^-1^). Mitochondria from bovine heart (5 µg), HEK293T and KRAS^G12V^/MYC (25 µg) were resuspended in 25 mM potassium-phosphate buffer pH 7.5 and supplemented with 500 µM KCN, 100 µM EDTA, 2 µM Rotenone and 100 µM DBQH_2_. The reaction was started using 75 µM oxidized cytochrome *c*. Complex III specific activity was determined by subtraction of the Antimycin A (2 µM)-sensitive activity.

#### Complex IV activity assay

Activity was determined by measuring cytochrome *c* oxidation (ε_550nm_ = 18.5 mM^-1*^cm^-1^). Mitochondria from bovine heart (5 µg), HEK293T and KRAS^G12V^/MYC (25 µg) mitochondria were resuspended in 25 mM potassium-phosphate buffer pH 7.0 and supplemented with 2 µM Antimycin A. The reaction was started using 75 µM of reduced cytochrome *c*. Complex IV specific activity was determined by subtraction of the KCN (500 µM)-sensitive activity.

#### Oncolines^®^ synergy profiling

To determine sublethal concentrations of (R)-GNE-140 and BMS-986205, dose-response curves were measured across nine concentrations (3.16 nM to 31.6 µM, √10 dilution steps) in duplicates. Cells were treated for 3 days in ATCC-recommended media, and proliferation was assessed. Cell stocks, maintained within 10 passages, were used to calculate doubling times from untreated cell growth at 0 and 72 hours. Assays were deemed invalid if doubling times deviated from specifications, prompting the use of a new cell stock. With the use of these data, identification of synergistic cell responses was conducted via dose-response profiling for the full dose-range of (R)-GNE-140 combined with a fixed concentration of BMS-986205 (10 µM) in the 102 Oncolines^®^ cell line panel. For specific cell lines (SR, CCRF-CEM, HT-1080, SU-DHL-1, PA-1, Jurkat E6.1, RKO), a lower dose of 3.16 µM BMS-986205 was used. Cell viability was determined via intracellular ATP quantification using ATP Lite™. For drug synergy calculation, the Bliss independence model was used ^104^. Bliss synergy score = E_AB_ – (E_A_ + E_B_ – (E_A_*E_B_)), where E_AB_ is the observed effect of the combination treatment and E_A_ + E_B_ – (E_A_ * E_B_) represents the predicted effect of the combination treatment using the measured effects of Drug A (E_A_) and Drug B (E_B_) monotreatment on cell-viability. Bioinformatic analyses were conducted by Oncolines^®^. Basal gene expression levels for 19,080 genes across 95 Oncolines^®^ cell lines were retrieved from the DepMap database (version 23Q4). Pearson correlations between the Bliss synergy scores of (R)-GNE-140 and BMS-986205 combination, as determined in Oncolines^®^ viability assays, and expression values of all genes in the corresponding cell lines were calculated. The significance of every calculated correlation was estimated by a p-value as determined in R. p-values were subjected to multiple testing correction using the Benjamini-Hochberg correction, based on the number of genes present in the various subsets. Pearson correlations with a false discovery rate <20% were considered significant. The mutation status of cell lines was determined using both public and proprietary datasets. Public data sources, including COSMIC Cancer Genome Project (version 80), were used to identify mutations, amplifications, and deletions in established cancer driver genes present in Oncolines^®^. To enhance accuracy, 23 cancer-associated genes were further validated via targeted and whole-exome sequencing directly from Oncolines^®^ cell lines. Genetic alterations were filtered based on their frequency in patient tumor samples from COSMIC, ensuring the exclusion of rare, non-cancer-relevant mutations. Gene Set Enrichment Analysis (GSEA) was performed using pre-ranked GSEA, as available in the *clusterProfiler* R package, with the full list of Person correlations as input using the KEGG database. Only gene sets containing a minimum number of 15 and a maximum number 500 genes were used in the analysis. The significance of a NES was determined by a permutation test. Analysis code and raw input files are available under 10.5281/zenodo.14591830.

#### Statistics and data reproducibility

No statistical methods were used to predetermine sample size. The experiments were conducted without randomization, and the investigators were not blinded to the allocation during both the experimental procedures and outcome assessment. Data analysis and plot generation were performed using GraphPad Prism 9, FlowJo, Fiji, ChemDraw, R and Microsoft Excel software. For the interaction of two variables, two-way ANOVA with multiple comparisons test (Tukey, Dunnett or Šidák), were performed as indicated. For time-course experiments, multiple two-sided t test was performed, which was followed by FDR correction (two-stage linear step-up procedure of Benjamini, Krieger, and Yekutieli). For the analysis of three or more groups, one-way ANOVA with multiple comparisons test (Tukey, Dunnett or Šidák), was performed as indicated. All Western blot, immunofluorescence-, brightfield- or electron microscopy experiments were replicated at least three times as indicated. Activity assays for respiratory chain complexes II – IV were performed as duplicates, as they were performed for three independent mitochondrial sources (bovine heart, KRAS^G12V^/MYC and HEK293T mitochondria). Experiments of Extended Data Fig. 5A regarding GNE+si-scramble and BMS+si-scramble were performed as duplicates as they closely recapitulated previously determined toxicity of GNE and BMS monotreatment.

## Reporting summary

Further information on research design is available in the Nature Research Reporting Summary linked to this article.

## Data availability

All data are provided in the Article, Source Data file and supplementary information.

## Acknowledgements

We thank Dr. Thomas Worzfeld for assisting with invasion assays, Dr. Natascha Sommer for her support with Seahorse experiments, and Dr. Guido Zaman (Oncolines) for valuable discussions. We also acknowledge expert technical assistance by Yvonne Horn and Markus Schwinn. We thank Natalie Weber for metabolite extraction and MS analysis. Our work is supported by funding from the German Research Foundation (DFG) GRK 2573/2 (RP10, Project 416910386), the Excellence Cluster CPI (Project 390649896), the EU (COST Action CA23119 Senescence2030) and the Forschungscampus Mittelhessen. H.F.F. was supported by the LOEWE Center Frankfurt Cancer Institute (FCI) funded by the Hessen State Ministry for Higher Education, Research and the Arts [III L 5 - 519/03/03.001 - (0015)] and the EU/EFPIA/OICR/McGill/KTH/Diamond Innovative Medicines Initiative 2 Joint Undertaking (EUbOPEN grant n° 875510). Disclaimer: This communication reflects the views of the authors, and the Joint Undertaking is not liable for any use that may be made of the information contained herein.

## Contributions

J.D., Conceptualization, Investigation, Data curation, Formal analysis, Writing-original draft; J.S., J.B., DvdB, V.M.B., M.H., C.L., B.N., A.N., T.S., J.T.-S., Investigation; H.F.F., M.P., A.C.-O., Methodology Formal analysis; U.G., R.S., Methodology; J.W., M.B., Formal analysis; J.G., S.K., Investigation, Data curation, Formal analysis; M.L.S., Supervision, Conceptualization, Funding acquisition, Writing-original draft.

## Ethics declarations

## Extended Data

**Extended Data Fig. 1.**
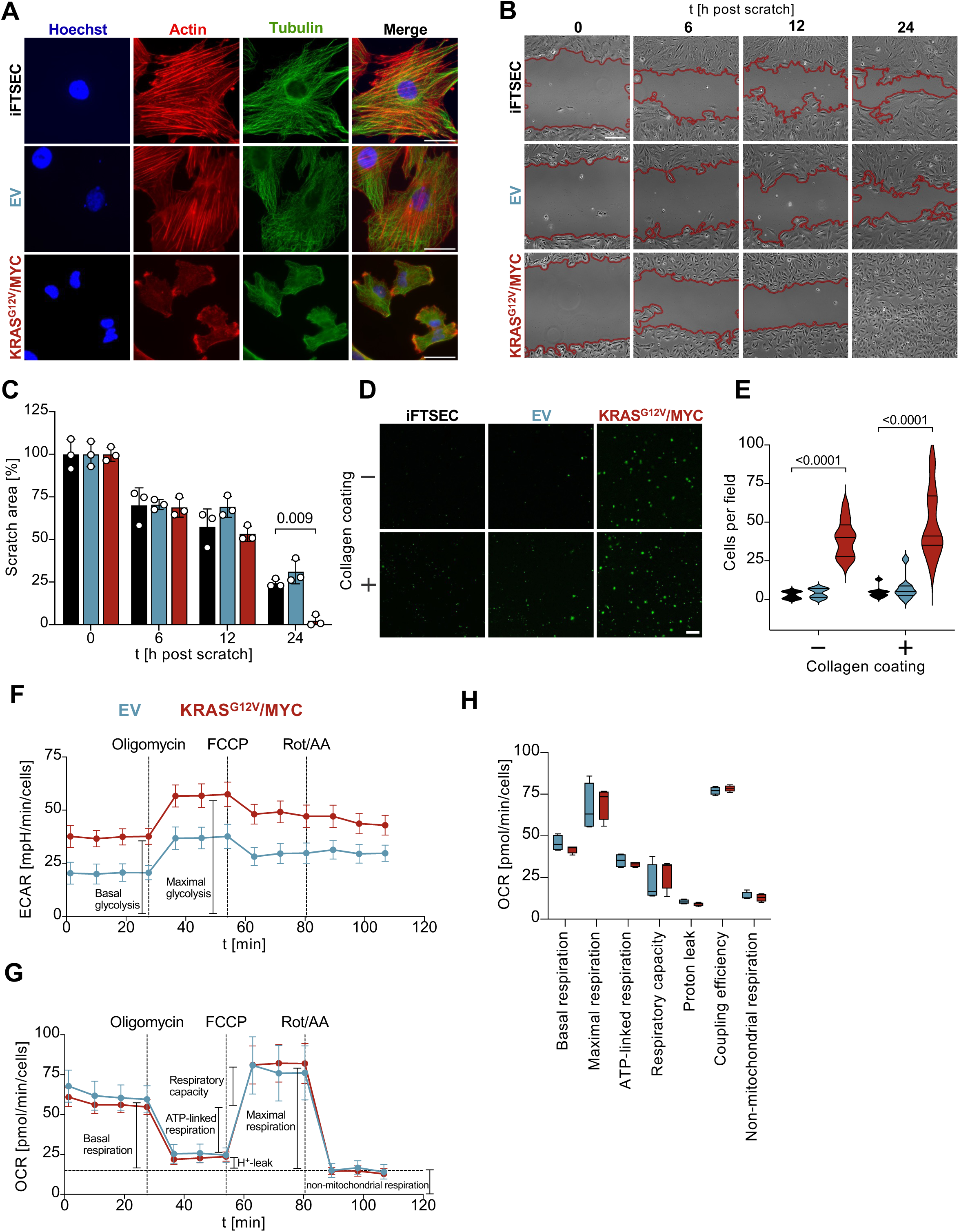
Characterization of oncogenic transformation of KRAS^G12V^/MYC cells. **(A)** iFTSEC, EV and KRAS^G12V^/MYC cells were stained for Actin and Tubulin by immunofluorescence microscopy. Representative examples are shown, the nuclear DNA was stained with Hoechst33342. Scale bar = 50 µm. **(B)** The indicated cells were grown to confluency and a scratch was made. Cells were further cultivated in medium containing only 1% (v/v) FCS to suppress proliferation. Closure of the scratch (indicated by red lines) was monitored over time. Scale bar = 200 µm. **(C)** Quantification of scratched areas. Shown are mean ± SD, *n* = 3, two-way ANOVA with Dunnett’s multiple comparisons test, the control was set to 100% **(D)** Transwell assay without collagen (monitoring migration) and collagen coating (monitoring invasion). Scale bar = 200 µm. **(E)** Quantification of migrated or invaded cells per field. Shown are violin plots, *n* = 3, two-way ANOVA with Dunnett’s multiple comparisons test. For the metabolic analysis displayed in the next subfigures, the indicated cell lines were analyzed by Seahorse metabolic flux analysis using the MitoStress test assay. At the indicated time points Oligomycin (2 µM), FCCP (2 µM) and Rotenone/Antimycin A (0.5 µM) were injected. **(F)** Analysis of glycolysis by measuring the extracellular acidification rate (ECAR). **(G)** Analysis of OXPHOS by measuring the oxygen consumption rate (OCR). **(H)** Quantification of different respiratory mechanisms. Shown are mean ± SD, *n* = 4, two-way ANOVA with Šídák’s multiple comparisons test.

**Extended Data Fig. 2.**
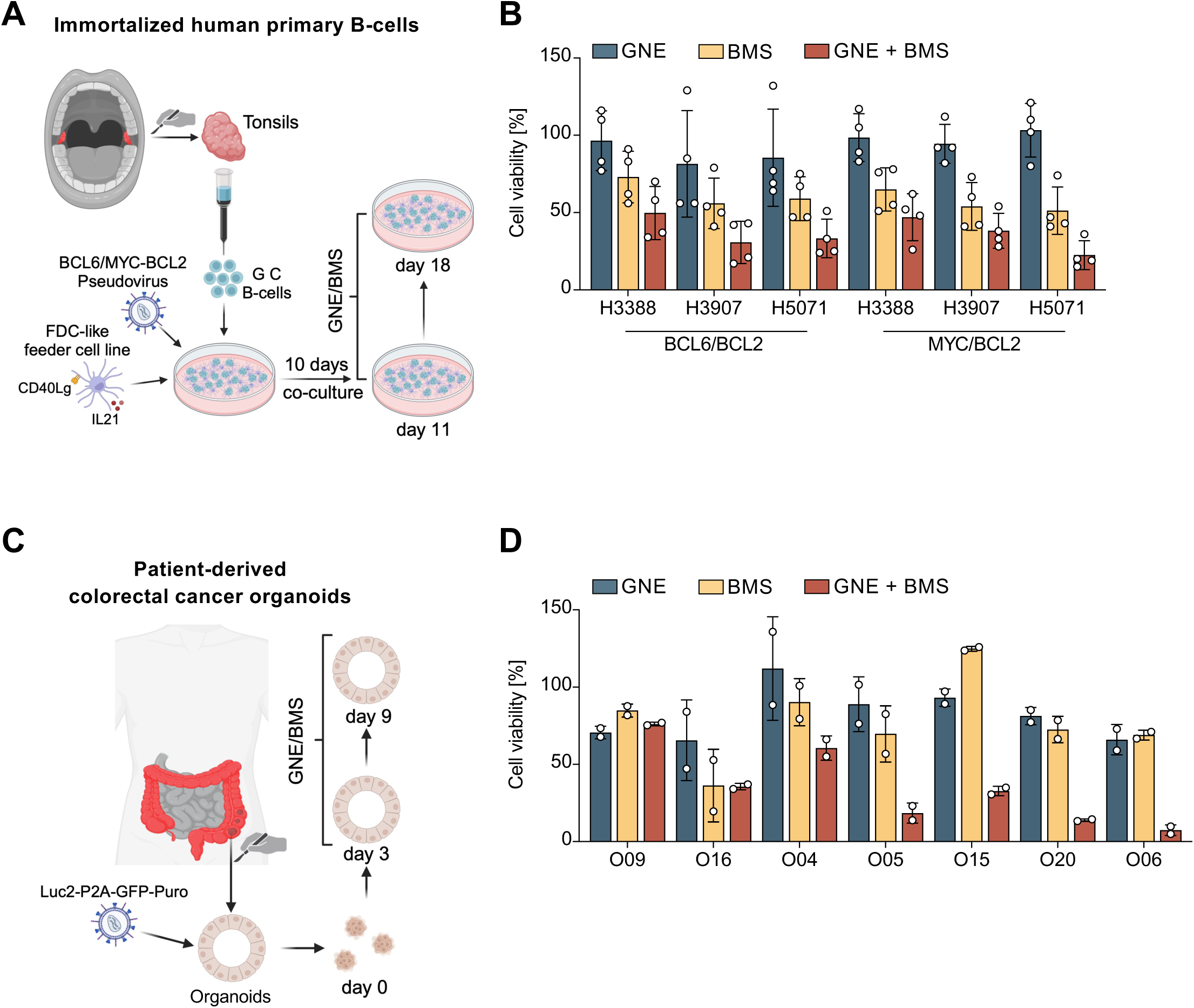
Impact of GNE/BMS treatment on human cell models. **(A)** Primary human germinal center B cells were isolated from tonsils, immortalized by viral delivery of BCL2 with BCL6 or MYC and further grown on feeder cells, as schematically shown. **(B)** Cells were sub-cultured for 10 days on irradiated FDC-like feeder cells prior treatment with sublethal concentrations of GNE and/or BMS for 7 days (suppl. Table 2). Cell number was determined by flow-cytometry using lymphocyte gating. Shown are mean ± SD, *n* = 4. **(C)** Colorectal cancer cells were collected, virally transduced to express a Luciferase-P2A-GFP reporter gene and grown to organoids. Following mono- and combination treatment with sublethal concentrations of GNE and BMS for 6 days (concentrations listed in suppl. Table 2), with a repeated treatment after 3 days, viability was assessed using the ONE-GloEX assays. **(D)** Results from the experiments schematically displayed in (C) are shown. Shown are mean ± SD, *n* = 2, two-way ANOVA with Šídák’s multiple comparisons test. All values were normalized to their respective DMSO vehicle control.

**Extended Data Fig. 3.**
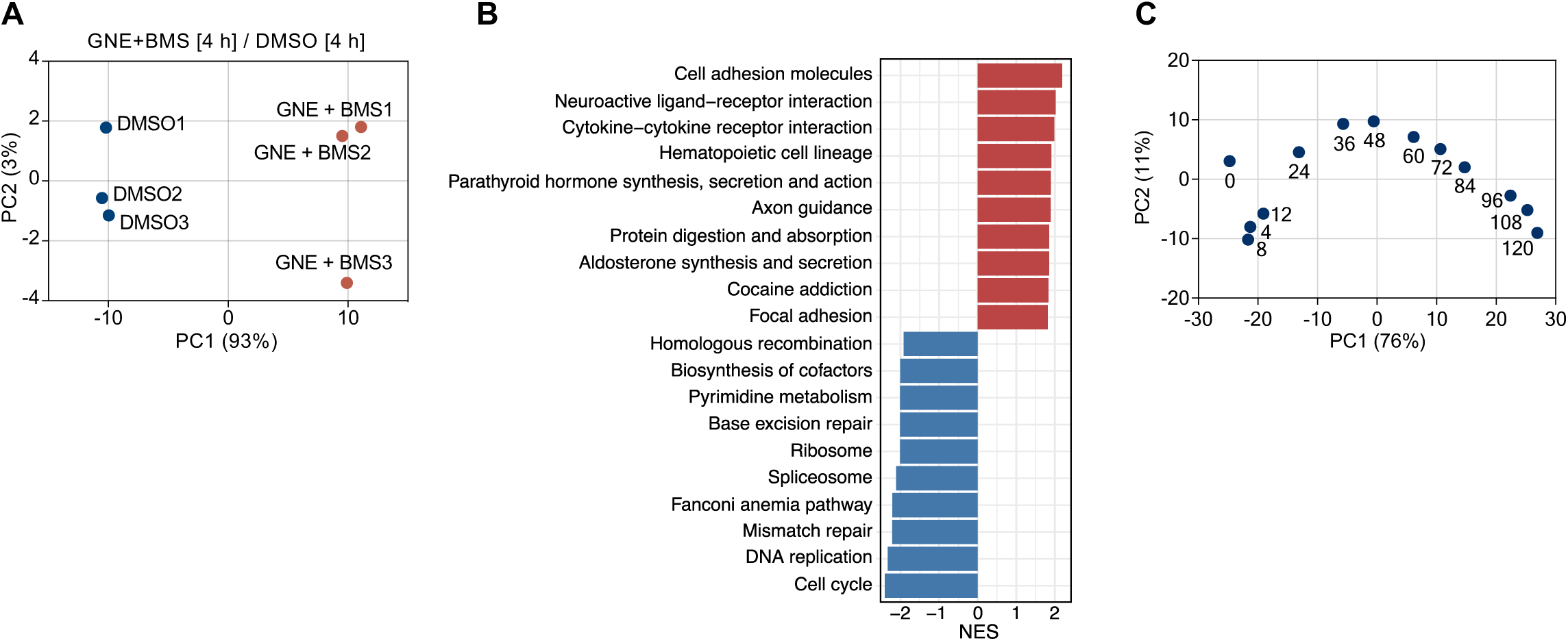
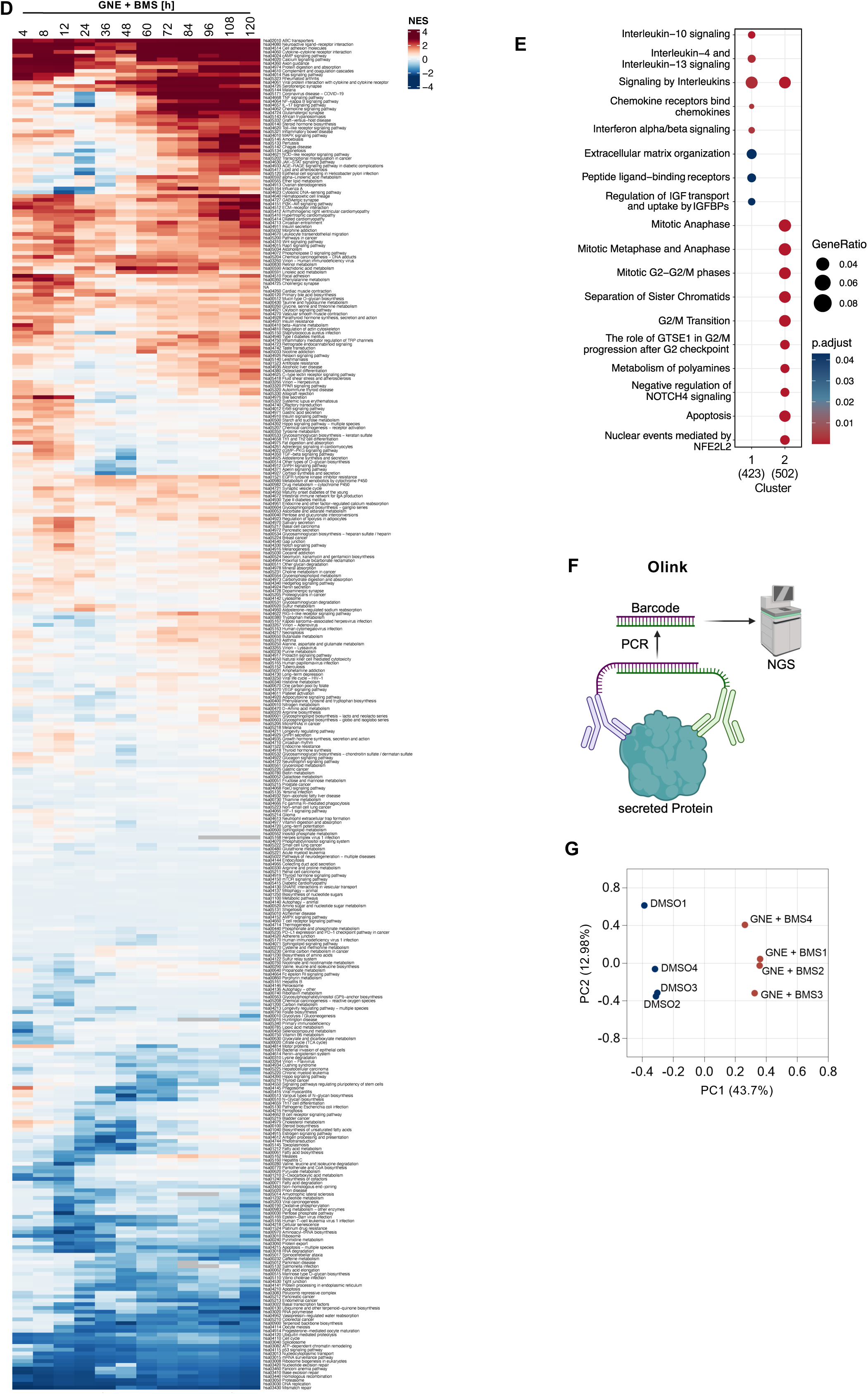
Characterization of GNE/BMS-mediated effects on RNA expression and protein secretion. **(A)** KRAS^G12V^/MYC cells were treated for 4 hours with GNE/BMS or vehicle (DMSO). A principal component (PC) analysis of RNA-seq data shows the first two components, samples are represented by dots, *n* = 3. **(B)** GSEA using the KEGG database of differentially expressed genes (Cut-off: log_2_FC ≥ 1, ≤ −1; FDR ≤ 0.05). **(C)** PC analysis of the longitudinal RNA-seq experiment over 120 hours post GNE/BMS treatment (Fig. 3B). **(D)** Heatmap displaying all KEGG pathways and their respective normalized enrichment score (NES) values over time. **(E)** GSEA of cluster 1 and 2 (Fig. 3B) using the Reactome database. **(F)** Schematic display of the workflow for the Olink^®^ analysis of secreted proteins. **(G)** KRAS^G12V^/MYC cells were treated for 4 days with GNE/BMS or DMSO. Supernatants were analyzed for secreted proteins using Olink^®^, a principal component analysis is shown, *n* = 4.

**Extended Data Fig. 4.**
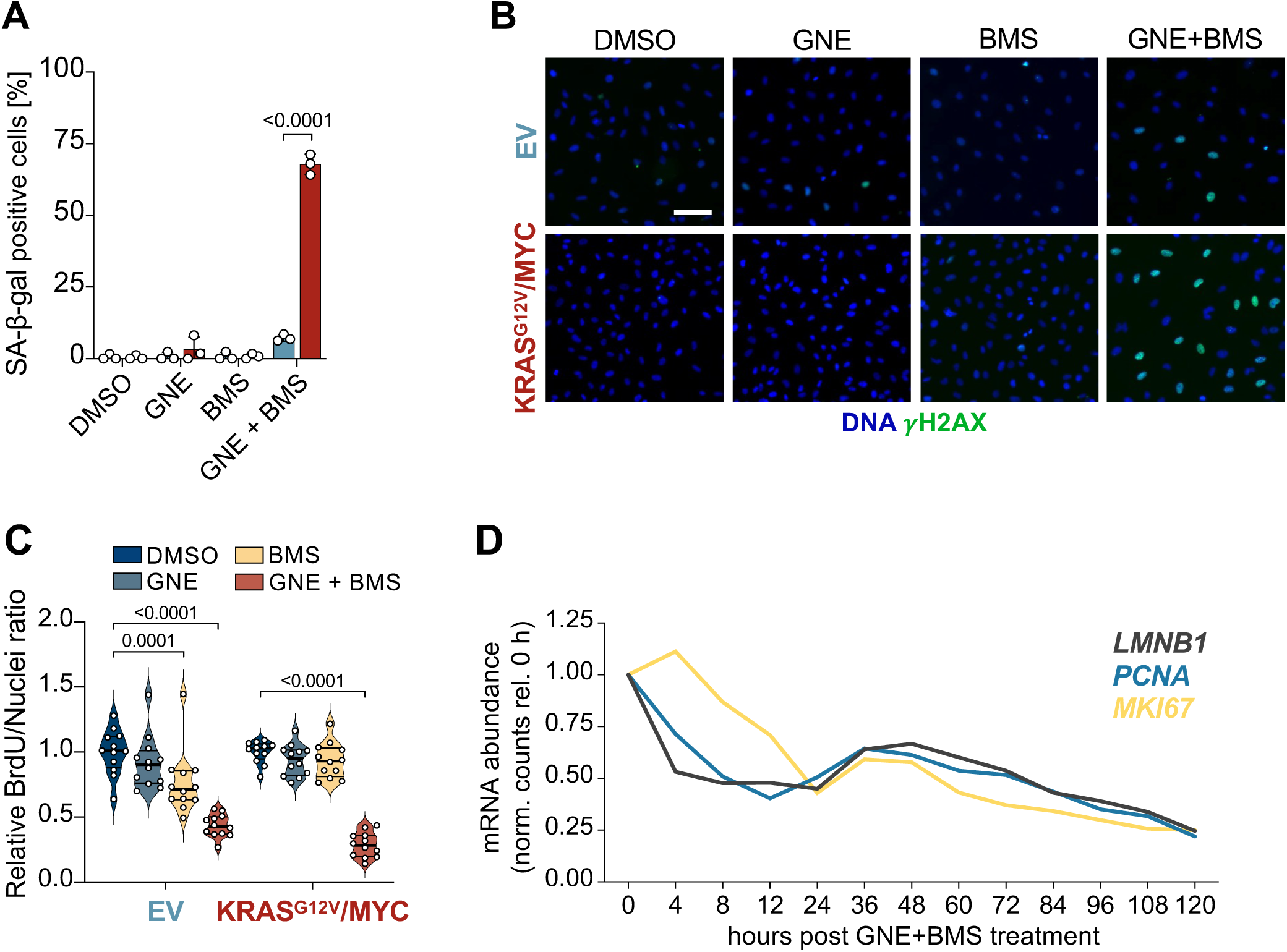
The combination of GNE and BMS induces cancer cell-specific senescence. **(A)** EV and KRAS^G12V^/MYC cell lines were analyzed for their SA-ß-gal activity after 3 days of GNE and BMS mono- or combination treatment. SA-ß-gal positive cells were manually counted and normalized to the total cell number per image, *n* = 3, two-way ANOVA with Šídák’s multiple comparisons test. **(B)** Representative γH2AX immunofluorescence images of EV control and KRAS^G12V^/MYC cancer cell lines after 2 days of GNE and BMS mono- or combination treatment. Scale bar = 100 µm. **(C)** Quantification of BrdU positive cells using immunofluorescence imaging after 2 days of GNE and BMS mono- or combination treatment. Shown are violin plots, *n* = 4, two-way ANOVA with two-stage linear step-up procedure of Benjamini, Krieger and Yekutieli. **(D)** Expression changes (Deseq2 normalization) of the indicated senescence-associated transcripts *MKI67*, *PCNA* and *LMNB1* (encoding Ki67, PCNA and Lamin B1) were derived from the RNA-seq data and normalized to the expression in untreated cells.

**Extended Data Fig. 5.**
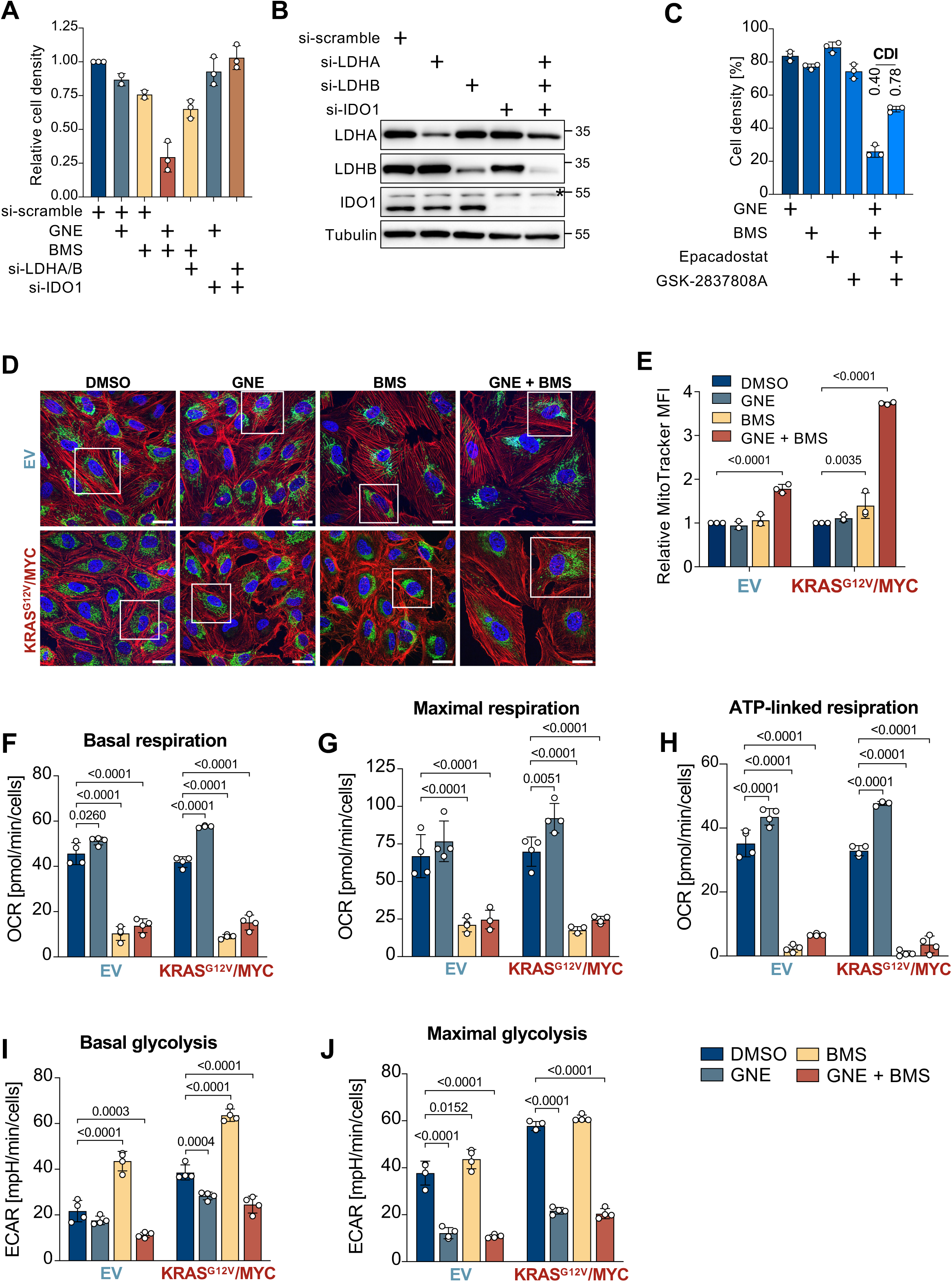
Mitochondrial effects of BMS. KRAS^G12V^/MYC cells were transfected with siRNAs targeting either LDHA/B or IDO1 alone or combined with GNE and/or BMS treatment as shown. Transfection was repeated after 2 days. **(A)** After 4 days one half of the cells was analyzed for effects on cell density, values were normalized to si-scramble, shown are mean ± SD, *n* = 3. **(B)** The other half of the siRNA-treated cells was analyzed by Western blotting for knock-down, the asterisk indicates a non-specific band, the positions of molecular weight markers are indicated. **(C)** Cells were treated for 3 days with Epacadostat (IDO1 inhibitor, 40 µM), GSK-2837808 (LDHA/B inhibitor, 35 µM) alone or in combination with GNE or BMS, as shown. Cell viability was determined, shown are mean ± SD, *n* = 3. The Coefficient of Drug Interaction (CDI) for GNE and BMS as well as Epacadostat and GSK-2837808 was calculated and indicated above respective bar plots. **(D)** COX8-GFP EV and KRAS^G12V^/MYC cells were treated for 2 days with the indicated conditions and stained with Hoechst33342 (nuclear DNA) and Phalloidin (Actin-filaments) following immunofluorescence analysis. White rectangular selections are highlighting the cells used for mitochondrial high-resolution 3D-rendering (Fig. 5A). **(E)** Cells treated with GNE and/or BMS for 2 days, loaded with MitoTracker Deep Red FM (200 nM for 15 minutes) and subsequently analyzed using flow-cytometry. Shown are median fluorescence intensities (MFI), *n* = 3, two-way ANOVA with Dunnet’s multiple comparisons test. The next subfigures show seahorse metabolic flux analyses of empty vector and KRAS^G12V^/MYC cell lines using the MitoStress Test assay after 4 hours of GNE and BMS mono- and combination treatment using Oligomycin (2 µM), FCCP (2 µM) and Rotenone/Antimycin A (0.5 µM). **(F-H)** Analysis of basal, maximal and ATP-linked respiration. **(I, J)** Quantification of basal and maximal glycolysis. Shown are mean ± SD, *n* = 4, two-way ANOVA with Šídák’s multiple comparisons test.

