## Supplemental Table 1 for "Synergistic targeting of cancer cells through simultaneous inhibition of key metabolic enzymes"

**Figure 1E: Inhibitors used in synthetic lethal screen**

| Name | Target | Sublethal concentration | Source | Cat # |
| --- | --- | --- | --- | --- |
| (±)-3-Methyl-2-  oxovaleric acid | KGDHC | 1 mM | Sigma | K7125 |
| (R)-GNE-140 | LDHA/B | 7.5 µM | MedChemExpr | HY-100742A |
| AZD-3965 | MCT1 | 5 µM | SellekChem | S7339 |
| BCH | LAT1 | 10 µM | Tocris | 5027 |
| BMS-986205 | IDO1 | 6 µM | MedChemExpr | HY-101560 |
| CB-839 | GLS | 10 µM | SellekChem | S7655 |
| Cytochalasin B | GLUT | 0.2 µM | Santa Cruz | sc-3519 |
| Dichloracetate | PDK | 1 mM | Sigma | 347795 |
| Eflornithin | ODC | 60 µM | Tocris | 2761 |
| Etomoxir | CPT1 | 50 µM | Sigma | S8244 |
| H3B-120 | CPS1 | 20 µM | Sigma | SML3007 |
| Indoximod | IDO/TDO | 2 µM | SelleckChem | S7756 |
| Metformin | Complex I | 10 mM | Invivogen | NC2257632 |
| PFK15 | PFKFB3 | 0.75 µM | Sigma | S7289 |
| TVB-2640 | FASN | 0.3 µM | Sigma | S9714 |
| V-9302 | ASCT2 | 3 µM | MedChemExpr | HY-112683 |
| α-cyano-4-hydroxycinnamic acid | MCT1 | 1 mM | Sigma | C2020 |
