## Supplemental Table 2 for "Synergistic targeting of cancer cells through simultaneous inhibition of key metabolic enzymes"

**Figure 2A: Sublethal concentrations of GNE and BMS for different cell lines**

| Cell line | RRID | Species | (R)-GNE-140 [µM] | BMS-986205 [µM] |
| --- | --- | --- | --- | --- |
| FS4-LTM  (from FS4) | CVCL_0266 | Human | 14 | 4 |
| HEK293T | CVCL_0063 | Human | 30 | 2.5 |
| HCT-116 | CVCL_0291 | Human | 20 | 8 |
| HeLa | CVCL_0030 | Human | 15 | 10 |
| HT-29 | CVCL_0320 | Human | 20 | 6 |
| iFTSEC (hTERT FT240) | CVCL_UH60 | Human | 7.5 | 6 |
| iFTSEC_EV | This study | Human | 7.5 | 6 |
| iFTSEC_KRAS^G12V^/MYC | This study | Human | 7.5 | 6 |
| KPC | CVCL_A9ZK | Mouse | 20 | 9 |
| LN229 | CVCL_0393 | Human | 20 | 20 |
| MCF-7 | CVCL_0031 | Human | 15 | 6 |
| mPSC4 | N/A | Mouse | 10 | 1 |
| OVCAR-4 | CVCL_1627 | Human | 7.5 | 6 |
| Panc^OVA^  (from Panc02) | CVCL_D627 | Mouse | 24 | 9 |
| S2-007 | CVCL_B279 | Human | 26 | 6 |
| PC9 | CVCL_B260 | Human | 15 | 8 |
| RPE-1 | CVCL_4388 | Human | 8 | 15 |
| SW620 | CVCL_0547 | Human | 10 | 6 |
| T110299 | N/A | Mouse | 26 | 10 |
| U2OS | CVCL_0042 | Human | 30 | 10 |

**Figure 2B: Sublethal concentrations of GNE and BMS for B cells**

| Cell line | (R)-GNE-140 [µM] | BMS-986205 [µM] |
| --- | --- | --- |
| MYC/BCL2 H5071 | 30 | 6 |
| MYC/BCL2 H3388 | 20 | 3 |
| MYC/BCL2 H3907 | 20 | 4 |
| BCL6/BCL2 H5071 | 20 | 4.5 |
| BCL6/BCL2 H3388 | 15 | 3 |
| BCL6/BCL2 H3907 | 20 | 4.5 |

**Figure 2D: Sublethal concentrations of GNE and BMS for patient-derived colorectal organoids**

| Patient organoid ID | (R)-GNE-140 [µM] | BMS-986205 [µM] |
| --- | --- | --- |
| O06 | 50 | 10.8 |
| O20 | 23.2 | 10.8 |
| O15 | 23.2 | 10.8 |
| O05 | 23.2 | 5 |
| O04 | 50 | 10.8 |
| O16 | 23.2 | 2.3 |
| O09 | 23.2 | 10.8 |
