## Supplemental Table 3 for "Synergistic targeting of cancer cells through simultaneous inhibition of key metabolic enzymes"

**Figure 8C**

| ANOVA results table for genetic transformation in 38 cancer genes and their statistical association with treatment synergy. Only mutations were considered that appeared at least 3 times in the Oncolines® cancer cell line panel. | | | |
| --- | --- | --- | --- |
| **Gene** | **Synergy shift** | **p-value** | **Adjusted p-value** |
| RB1 | 28.40 | 4.42e-04 | 1.68e-02 |
| EP300 | 23.08 | 4.06e-03 | 7.71e-02 |
| LRP1B | 18.16 | 6.76e-03 | 8.57e-02 |
| NSD1 | -16.47 | 2.50e-02 | 2.38e-01 |
| XIRP2 | -6.98 | 4.41e-02 | 3.35e-01 |
| BRAF | 15.31 | 6.61e-02 | 4.13e-01 |
| EGFR | -10.24 | 7.62e-02 | 4.13e-01 |
| ABL.driven | 17.54 | 8.69e-02 | 4.13e-01 |
| ARID1A | -8.23 | 1.25e-01 | 4.83e-01 |
| KRAS | 3.34 | 1.57e-01 | 4.83e-01 |
| FBXW7 | -18.69 | 1.66e-01 | 4.83e-01 |
| CTNNB1 | -11.99 | 1.89e-01 | 4.83e-01 |
| APC | 15.34 | 2.05e-01 | 4.83e-01 |
| CCND1 | -3.61 | 2.12e-01 | 4.83e-01 |
| STK11 | 0.25 | 2.15e-01 | 4.83e-01 |
| SMAD4 | -3.72 | 2.17e-01 | 4.83e-01 |
| NF1 | 4.49 | 2.21e-01 | 4.83e-01 |
| BRCA2 | 6.87 | 2.29e-01 | 4.83e-01 |
| CCNE1 | -15.56 | 2.42e-01 | 4.83e-01 |
| CDKN2A | 0.78 | 3.40e-01 | 6.03e-01 |
| ZFHX3 | 0.43 | 3.47e-01 | 6.03e-01 |
| SETD2 | -5.16 | 3.55e-01 | 6.03e-01 |
| NCOR1 | 20.18 | 3.65e-01 | 6.03e-01 |
| ATM | -3.94 | 4.07e-01 | 6.31e-01 |
| PIK3R1 | 3.96 | 4.15e-01 | 6.31e-01 |
| ERBB2 | -8.05 | 4.97e-01 | 7.26e-01 |
| SMARCA4 | 2.56 | 5.76e-01 | 7.93e-01 |
| CHD4 | 24.74 | 5.84e-01 | 7.93e-01 |
| MYC | -5.40 | 6.28e-01 | 8.23e-01 |
| PTEN | -6.31 | 7.17e-01 | 8.83e-01 |
| NOTCH1 | 0.10 | 7.42e-01 | 8.83e-01 |
| SPEN | 8.01 | 7.44e-01 | 8.83e-01 |
| NRAS | -7.76 | 8.16e-01 | 9.18e-01 |
| FAT1 | -7.15 | 8.22e-01 | 9.18e-01 |
| TP53 | 1.96 | 8.56e-01 | 9.29e-01 |
| PBRM1 | -9.10 | 9.06e-01 | 9.55e-01 |
| CREBBP | -0.98 | 9.30e-01 | 9.55e-01 |
| PIK3CA | 7.14 | 9.78e-01 | 9.78e-01 |
