## Supplemental Table 4 for "Synergistic targeting of cancer cells through simultaneous inhibition of key metabolic enzymes"

**Figure 8C**

| Mutations and copy number variations for the cancer genes showing significant enrichment | | | | |
| --- | --- | --- | --- | --- |
| **Gene** | **Cell line** | **Protein change** | **Mutation type** | **Mutation zygosity** |
| EP300 | BxPC-3 | p.R397* | Substitution - Nonsense | het |
| EP300 | DLD-1 | p.R838C | Substitution - Missense | het |
| EP300 | DLD-1 | p.E1014* | Substitution - Nonsense | het |
| EP300 | HCT 116 | p.M1470fs*26 | Deletion - Frameshift | het |
| EP300 | HCT 116 | p.N1700fs*9 | Deletion - Frameshift | het |
| EP300 | HCT 15 | p.E1014* | Substitution - Nonsense | hom |
| EP300 | HT | p.N1547fs*17 | Deletion - Frameshift | het |
| EP300 | LS411N | p.H2324fs*55 | Insertion - Frameshift | het |
| EP300 | MCF-7 | p.R1356* | Substitution - Nonsense | het |
| EP300 | MOLT-4 | p.M1470fs*26 | Deletion - Frameshift | het |
| EP300 | RKO | p.K292fs*25 | Deletion - Frameshift | het |
| EP300 | RKO | p.M1470fs*26 | Deletion - Frameshift | het |
| EP300 | RL | p.Y1414C | Substitution - Missense | het |
| EP300 | RL | p.E1011* | Substitution - Nonsense | het |
| EP300 | RL95-2 | p.H2324fs*55 | Insertion - Frameshift | het |
| EP300 | SU-DHL-6 | p.R1627W | Substitution - Missense | het |
| EP300 | SW48 | p.M1470fs*26 | Deletion - Frameshift | het |
| EP300 | SW620 | p.P1440L | Substitution - Missense | het |
| EP300 | T24 | p.C1201Y | Substitution - Missense | hom |
| LRP1B | Hs 766T |  | Gene deletion |  |
| LRP1B | NCI-H460 |  | Gene deletion |  |
| LRP1B | OVCAR-3 |  | Gene deletion |  |
| LRP1B | U2OS |  | Gene deletion |  |
| RB1 | 5637 | p.Y325* | Substitution - Nonsense | hom |
| RB1 | BT-20 | p.P515L | Substitution - Missense | het |
| RB1 | BT-549 |  | Gene deletion |  |
| RB1 | DU145 | p.K715* | Substitution - Nonsense | hom |
| RB1 | DU4475 |  | Gene deletion |  |
| RB1 | TCCSUP |  | Gene deletion |  |
